# The GPI sidechain of *Toxoplasma gondii* prevents parasite pathogenesis

**DOI:** 10.1101/2024.02.21.581431

**Authors:** Julia A Alvarez, Elisabet Gas-Pascual, Sahil Malhi, Ferdinand Ngale Njume, Juan C Sánchez-Arcila, Hanke van der Wel, Yanlin Zhao, Gabriella Ceron, Jasmine Posada, Scott P Souza, George S Yap, Christopher M West, Kirk D C Jensen

## Abstract

Glycosylphosphatidylinositols (GPIs) are highly conserved anchors for eukaryotic cell surface proteins. The apicomplexan parasite, *Toxoplasma gondii*, is a widespread intracellular parasite of warm-blooded animals whose plasma membrane is covered with GPI-anchored proteins, and free GPIs called GIPLs. While the glycan portion is conserved, species differ in sidechains added to the triple mannose core. The functional significance of the Glcα1,4GalNAcβ1-sidechain reported in *Toxoplasma gondii* has remained largely unknown without an understanding of its biosynthesis. Here we identify and disrupt two glycosyltransferase genes and confirm their respective roles by serology and mass spectrometry. Parasites lacking the sidechain on account of deletion of the first glycosyltransferase, PIGJ, exhibit increased virulence during primary and secondary infections, suggesting it is an important pathogenesis factor. Cytokine responses, antibody recognition of GPI-anchored SAGs, and complement binding to PIGJ mutants are intact. In contrast, the scavenger receptor CD36 shows enhanced binding to PIGJ mutants, potentially explaining a subtle tropism for macrophages detected early in infection. Galectin-3, which bind GIPLs, exhibits a slight enhancement of binding to PIGJ mutants, and the protection of galectin-3 knockout mice from lethality suggests that *Δpigj* parasite virulence in this context is sidechain dependent. Parasite numbers are not affected by *Δpigj* early in the infection in wildtype mice, suggesting a breakdown of tolerance. However, increased tissue cysts in the brains of mice infected with *Δpigj* parasites indicate an advantage over wildtype strains. Thus, the GPI sidechain of *T. gondii* plays a crucial and diverse role in regulating disease outcome in the infected host.

**Summary:** The functional significance of sidechain modifications to the GPI anchor is yet to be determined because the glycosyltransferases responsible for these modifications have not been identified. Here we present identification and characterization of both *T*. *gondii* GPI sidechain-modifying glycosyltransferases. Removal of the glycosyltransferase that adds the first GalNAc to the sidechain results in parasites without a sidechain on the GPI, and increased parasite virulence. Loss of the second glycosyltransferase results in a sidechain with GalNAc alone, and no glucose added, and has negligible effect on parasite virulence. This indicates GPI sidechains as fundamental to host-parasite interactions.

## Introduction

Protozoan parasites are widespread and cause prominent diseases including malaria, leishmaniasis, chagas, and toxoplasmosis. One notable feature of protozoans is their extensive decoration of glycosylinositolphospholipids (GIPL) and GPI-anchored proteins (GPI-AP). The GPI was first discovered and characterized in trypanosomes, the protozoan parasite that causes African sleeping sickness in humans (1). Since its discovery, it has been shown to have a conserved core structure across the eukaryotic kingdom; EtNP-6Manα1-2Manα1-6Manα1-4GlcN1-6*myo*-inositol-phospholipid (where EtNP, Man, and GlcN are ethanolamine phosphate, mannose, and glucosamine, respectively) (1,2). Apicomplexan parasites use GPI-APs to assist attachment to and cellular invasion of host cells (3) and as such, the core GPI synthetic pathway enzymes are each essential gene products required for *Toxoplasma gondii* (*T. gondii)* and *Plasmodium sp*. survival and intracellular infection (4,5).

In the case of *T. gondii*, a widespread apicomplexan parasite of warm-blooded animals and humans, cellular attachment and invasion are mediated by a large family of *T. gondii* GPI-APs called surface antigen glycoprotein (SAG)-related super family (SRS) (6,7). It is known that these dominant SAG antigens are targeted by *T. gondii*-specific antibodies following infection (8), but also play important roles in virulence. For example, SAG1, the dominant antigen in the parasite’s lytic stage has been shown to promote small intestinal ileitis in mice (9), and promotes parasite survival *in vivo* (10) and in activated macrophages, by an unknown mechanism (11). Another GPI-AP of *T. gondii*, SRS35 (also known as p18 or SAG4), has been shown to promote mouse macrophage invasion and virulence of the parasite (12). In contrast, overexpression of the GPI-AP SRS29C (p35) quells *T. gondii* virulence and promotes mouse survival to an otherwise lethal infection (6).

While many GPI-APs have been considered for their role in parasite virulence, the GPI that anchors them has long been shown to be immunogenic in *T. gondii* and other parasites. For instance, *T. cruzi* GIPLs are potent activators of TLR2 (13). Similarly, GIPLs of *P. falciparum* are thought to be major pathogenesis factors, and are shown to activate TLR2 and TLR4 (14), which in turn causes lethal inflammatory responses (15). The GIPL of *T. gondii* activates TLR2 and TLR4 (16). *T. gondii* and *Plasmodium sp.* GIPLs are also robustly targeted by antibodies during the early stages of infection in both humans and in mice (17–25). Collectively, these data demonstrate that parasitic GPIs illicit robust innate and adaptive immune responses during infection.

Whereas the core GPI structure is conserved across the eukaryotic kingdom, species differ in how their GPI is modified through the attachment of additional sugars and other moieties to the mannose core, called sidechain modifications (26–28) (**Fig S1**). The significance of these sidechains and why species possess structural diversity in their GPI modifications is unclear because the responsible GPI glycosyltransferases (GT) have remained unidentified in most species. However in humans, the glycosyltransferase PGAP4 was recently determined to be the GT responsible for adding D-GalNAc to the first mannose of the human GPI, the first subunit of the sidechain (29). PGAP4 is widely conserved among eukaryotic species that are similarly substituted, including *Caenorhabditis elegans*. In the case of *T. gondii,* the GPI sidechain occurs on the same first mannose in two glycoforms: one consisting of a single identically linked D-GalNAc, and the other in which the GalNAc is extended by an α4-linked D-Glc (17). Interestingly, however, PGAP4 is not conserved in *T. gondii* despite sharing the same GalNAc branch at the first mannose (**Fig S1**) (29). When the PGAP4 homolog was deleted in mice the deficiency resulted in elevated blood alkaline phosphatase levels, impaired bone formation, decreased locomotion, impaired memory, and more enhanced vulnerability to prion diseases (30). These results point to GPI sidechains having far-reaching and important roles in mammals.

In the context of eukaryotic microbes, even less is known regarding the functional significance of the GPI sidechain. For example, in *Plasmodium* and fungi there is a fourth mannose added as a sidechain to the third mannose of the core GPI by the *SMP3* glycosyltransferase (**Fig S1**). Removal of *SMP3* in *Candida albicans* interferes with fungal viability *in vitro* (31). In *P. falciparum* which has a fourth mannose GPI sidechain, but lacks a homolog to *SMP3*, its GT *PIGB* that is responsible for adding the third mannose to the GPI core can add the fourth mannose when complemented into *C. albican Δsmp3* mutants. Whether *P. falciparum PIGB* encodes the GT responsible for adding the mannose GPI sidechain is uncertain because *Δpigb* parasites fail to grow *in vitro* (32). Multiple GPI sidechain GT’s have been identified in *Trypanosoma brucei* (*T. brucei*) parasites which exhibit multiple elaborately branched sidechains emanating from the second mannose of its GPI core. The *T. brucei* GT *TbGT8*, which adds the first branch point of one of the sidechain branches, was shown to be nonessential in bloodstream- and procyclic-form stages of the parasite (33). *TbGT3* is expressed in the bloodstream and procyclic stages of *T. brucei*, and mutants remained infectious to tsetse flies, though the Gal addition to the non-reducing-terminal GlcNAc residue of the GPI sidechain was lost (34). Loss of *TbGT10* resulted in a minor fitness cost evidenced by reduced growth *in vitro*, likely a consequence of impaired carbohydrate synthesis, but *TbGT10* was responsible for adding all β6 linkages to the GPI sidechain, resulting in 10 less sugars in *TbGT10* null parasites (35). Although removal of the aforementioned *T. brucei* GT’s reduces some of the GPI sidechain branching complexity, none of these GT mutants lack a GPI sidechain, suggesting functional redundancy may exist between different GPI glycoforms to support *T. brucei* fitness. Whether expression of specific glycoforms of the GPI are required for *in vivo* fitness of eukaryotic microbes is largely unknown.

While there have been several eukaryotic GPI sidechain modifying glycosyltransferases identified to date, none have addressed whether lacking a GPI sidechain entirely or whether specific glycoforms are required for microbial pathogenicity. Therefore, we sought to address these questions in *T. gondii* and test whether specific GPI glycoforms impact parasite virulence and immune recognition. Here we identify the two sidechain-modifying glycosyltransferases in *T. gondii* and report that loss of the GPI sidechain promotes virulence. Whereas antibody recognition of parasite GPI-APs and inflammatory cytokines appear intact to GPI sidechain null parasites, we present indirect evidence for galectin-3 and the scavenger receptor, CD36, in mediating certain phenotypes associated with sidechain deficiency in *T. gondii* strains.

## MATERIALS & METHODS

### Parasite strains and cell lines

Human foreskin fibroblasts (HFFs) monolayers were grown in DMEM GlutaMAX^TM^ (4.5 g/L D-glucose) (Life Technologies #10-566-024) supplemented with 2 mM L-glutamine, 20% heat inactivated (HI) fetal bovine serum (FBS) (Omega Scientific), 1% penicillin-streptomycin (Life Technologies, #15140122), and 0.2% gentamycin (Life Technologies #15-710-072). Mouse Embryonic Fibroblasts (MEFs) were grown in DMEM GlutaMAX^TM^ (4.5 g/L D-glucose) supplemented with 10% HI FBS, 20mM HEPES, 1% penicillin-streptomycin, and 0.2% gentamycin. *Toxoplasma gondii* strains were passaged in HFFs in “Toxo medium” (4.5 g/L D-glucose, L-glutamine in DMEM GlutaMAX^TM^ supplemented with 1% HI FBS and 1% penicillin-streptomycin). The following clonal strains were used (clonal types are indicated in parentheses): RH *Δku80 Δhxgprt* (type I), RH *GFP:cLUC* (type I) (clone 1-1), RH *Δku80 Δhxgprt Δpigj::HXGPRT* (type I) (clone C12), RH *Δku80 Δhxgprt Δpige::DHFR-TS* (type I) (36), GT1 (type I), GT1 *Δpigj:DHFR-TS* (type I) (clone B8), GT1 *Δpige::DHFR-TS* (type I) (clone C11), CEP *hxgprt*-(type III), CEP *hxgprt-GFP:cLUC* (type III), CEP *hxgprt*-*Δpigj:hxgprt GFP:cLUC* (type III) (clone D10), CEP *hxgprt*-*Δpigj::HXGPRT PIGJ-HA_3x_:DHFR-TS GFP:cLUC* (type III) (clone C2), CEP *hxgprt*-*Δpige::hxgprt GFP:cLUC* (type III) (clone D4). All strains, oligos and plasmids used for the generation of strains in this study can be found in **Table S1**.

### Generation of gene edited *T. gondii* strains

Cas-9 and single guide RNA (gRNA) expression plasmid, pSS013 (gift from Jeroen Saeij, University of California, Davis) was designed to target *Tg_207750* (Uniport S8FC68)*, PIGJ*. For *Tg_266320* (Uniprot S7VW57)*, PIGE* a modified dual-guide pU6-universal plasmid targeting two different exons within the gene was previously generated (36). Selectable markers were produced by PCR amplification. An amplicon of the pyrimethamine selectable cassette (*DHFR-TS)* was amplified from the pLoxp-DHFR-TS-mCherry plasmid (Addgene plasmid #70147) containing similar homology arms to the targeted protospacer sequences. The amplicon of *HXGPRT* was generated from the plasmid pTKO-att (a gift from Jeroen Saeij, University of California, Davis) with homology arms flanking the targeted protospacer sequences of the targeted gene. The expression Cas9/gRNA plasmid and amplicon were co-transfected via electroporation at a 5:1 ratio. Transfectants were selected and cloned in medium containing pyrimethamine (1 μM) or 50 μg/mL of MPA (Axxora) and 50 μg/mL of xanthine (Alfa Aesar) and screened for the disruption the locus by diagnostic PCR.

### Generation of the *PIGJ* complementation construct and complementation strain

The pLIC-HA-DHFR-TS (gift from Jeroen Saeij, University of California, Davis) plasmid was treated simultaneously with PacI (NEB #R0547S) and AvrII (NEB #R0174S) restriction endonucleases to allow for insertion of *TgVEG_PIGJ* by homology directed ligation. The full-length coding region with introns spliced out of *PIGJ* was amplified using the Q5 high fidelity polymerase (NEB, M0491) from a CEP cDNA library preparation using primers designed to contain 19 bp homology with the 3’ end of the *PIGJ* promoter sequence and 25 bp homology with end of the digested pLIC-HA plasmid that contains and provides a 3x HA tag, respectively. The CEP cDNA library was prepared from CEP total RNA preparation using the High-Capacity cDNA Reverse Transcription Kit (ThermoFisher, cat# 4368814) according to manufacturer’s instructions. For amplification of the *PIGJ* promoter region, a 1000 bp non-coding region upstream of the start ATG of *PIGJ* was amplified from CEP genomic DNA with Q5 high fidelity polymerase according to manufacturer’s protocol. The forward primer contained a 19 bp homology sequence to the end of the digested pLIC-HA plasmid and the reverse shared a 22 bp homology sequence with the 5’ end of the amplified PIGJ coding-cDNA sequence described above. The amplified promoter and coding *PIGJ* DNA sequence were first purified from 1% agarose gels using the ZymocleanTM Gel DNA recovery kit (Zymo Research, #D4007) and assembled in frame into the digested pLIC-HA -DHFR-TS plasmid in frame with the 3x HA tag using the NEBuilder HiFi DNA Assembly cloning Kit (NEB, cat# E5520) according to the manufacturer’s instructions. The CEP *hxgprt-Δpigj::HXGPRT GFP:cLUC* strain was transfected with the linearized pLIC-HA-DHFR-TS PIGJ-HA3x plasmid, grown in pyrimethamine selection medium and cloned by limiting dilution to generate the *CEP Δpigj::PIGJ-HA_3x_* complementation strain.

The HA tag was visualized by fluorescent microscopy. 8×10^5^ HFFs were plated on coverslips in 24-well plates overnight before being infected with 8×10^4^ parasites overnight. Cells were fixed with 3% formaldehyde and permeabilized with blocking buffer (1X PBS with 10% normal goat serum, and 0.01% saponin) and stained with rat anti-HA (1:50) (3F10, Sigma). Secondary goat anti-rat AF594 (Thermo Fisher cat#A11007) antibodies were used at (1:3000), and DAPI (1:10000), for detection and visualized using a fluorescent microscope (Nikon Eclipse T*i*-5).

### Mice

C57BL/6J (B6), A/J, and *Lgals3-/- (*B6.Cg-*Lgals3^tm1Poi^/*J) mice were purchased from Jackson Laboratories. *Tlr2/4* double knockout B6 mice (B6.129P2*-Tlr4^tm1Aki^* B6.129P2*-Tlr2^tm1Aki^*) were a generous gift from Dr. Greg Barton (UC Berkeley). Mice were maintained under specific pathogen free conditions at UC Merced. Female mice of 6-10 weeks of age were used for experiments unless otherwise stated.

### Ethics statement

Mouse work was performed in accordance with the National Institutes of Health Guide to the Care and Use of Laboratory Animals. Mouse protocols used have been reviewed and approved by UC Merced’s Committee on Institutional Animal Care and Use Committee (IACUC, #AUP-20-0015). UC Merced has an Animal Welfare Assurance filed with OLAW (A4561-01), is registered with USDA (93-R-0518), and the UC Merced Animal Care Program is AAALAC accredited (001318).

### Parasite infections and serotyping

Parasite injections were prepared by scraping T-25 flasks containing vacuolated HFFs and sequential syringe lysis first through a 25G needle followed by a 27G needle. The parasites were spun at 34 x g for 5 minutes to remove debris and the supernatant was transferred, followed by a spin at 611 x g and washing with sterile 1X PBS (Life Technologies, #10-010-049). For primary infections, mice were infected intraperitoneally (i.p.) with 104 tachyzoites of type III CEP *hxgprt-* parental and mutant strains. Parasite viability of the inoculum was determined by a plaque assay. In brief, 100 or 300 tachyzoites were plated in HFF monolayers grown in a 24-well plate and 4-6 days later were counted by microscopy (4x objective) (Nikon Eclipse Ti-5).

At 30 to 35 days after primary infection, 50μl of blood was harvested and collected in Eppendorf tubes containing 5μL 0.5 M EDTA and placed on ice. Blood was pelleted at 9391 x g for 5 minutes, and blood plasma was collected from the supernatant and stored at -80°C. To evaluate the seropositivity of the mice, HFFs were grown on coverslips and infected with green fluorescent protein (GFP)-expressing RH (1–1) overnight. 18 hrs later, cells were fixed with 3% formaldehyde in PBS, permeabilized with a permeabilization solution (3% bovine serum albumin fraction V (Omega, FB-11), 2% normal goat serum (Omega, NG-11) 0.2 M Triton X-100, 0.01% sodium azide), incubated with a 1:100 dilution of collected blood plasma for 2 hrs at room temperature, washed with 1X PBS, and detected with Alexa Fluor 594-labeled goat secondary antibodies specific for mouse IgG (Thermo Fisher, cat#A11032) in permeabilization solution. Seropositive parasites were observed by immunofluorescence microscopy.

For secondary infections, seropositive mice were infected i.p. with 5×10^4^ tachyzoites of type I RH *Δ*ku80 parental and mutant strains. Parasite viability of the inoculum was determined by plaque assay described above.

### Cyst enumeration using Dolichos-FITC

To quantify brain cysts, brains were dissected and placed in 10mLs of 1X PBS on ice. Brains were homogenized using a 10mL syringe with a 18G needle by extrusion several times through the needle. The homogenate was spun at 611 x g for 7 minutes and then resuspended in 1mL of 1X PBS. 100μL of the brain homogenate was fixed in ice cold methanol at a 1:10 dilution for 5 minutes, spun in a microcentrifuge at 5200 x g for 5 minutes and washed once and resuspended in 1mL 1X PBS. The fixed homogenate was stained with a 1:150 dilution of FITC-conjugated Dolichos biflorus agglutinin (Vector Laboratories cat# FL-1031) in 1X PBS and slowly rotated at 4°C overnight. Samples were washed twice with 1X PBS and resuspended in 1mL of 1X PBS. 50μL of stained homogenate was plated in a 96 well plate and four replicates of each sample were enumerated for cysts with an inverted fluorescence microscope using a 20x objective (Nikon Eclipse T*i*-5).

### Superinfections

Mice that survived 35 days of secondary infection were euthanized and brain homogenates were generated and resuspended in 1mL of 1X PBS as described above. 100μL of the homeogenate was cultured in HFFs and monitored for parasite growth. Once parasite growth was established, the media was switched to selection media containing MPA/xanthine for secondary infections to select against the primary infecting CEP *hxgprt-* strain, and for *HXGPRT* expressing secondary infection strain. Parasites that grew in selection media were also confirmed to be type I strains through restriction fragment length polymorphism analysis. In brief, isolated DNA from MPA/xanthine selected parasites was PCR amplified using GRA6 primers and the purified amplicon was fragmented with MseI (NEB cat# R0525M). Products were run on a gel to determine parasite type I or III based on unique fragmentation sizes indictive of the genotype.

### SDS-PAGE and immunoblotting for parasite lysate antigen

To generate parasite lysate antigens *T. gondii* was cultured in HFF monolayers and expanded to approximately 2×10^8^ parasites in a T175 flask. Parasites were syringe-lysed, washed with sterile 1X PBS, and pelleted at 611 x g for 7 minutes. The pelleted parasites were lysed with Lameli Buffer (0.0625M Tris Base, 0.07M SDS, 10% glycerol, 5% β-mercaptoethanol) and centrifuged at 14,000 x g for 20 minutes to remove large insoluble debris. The supernatant was aliquoted and stored at -80°C. Parasite lysates were separated via SDS-PAGE in 4-20% acrylamide hand cast gels before transfer to PVDF membrane. Membranes were blocked with 10% fortified bovine milk (Raley’s) dissolved in Tris-Buffered Saline with 0.1% Tween (TBS-T 0.1%) for 1-2 hrs at room temperature or overnight at 4°C. Blots were then probed with *T. gondii* GPI anchor glycoform-specific antibody clones T3 3F12 (BEI) which binds GalNAc GIPL, or T5 4E10 (a gift from Jean Francois Dubremetz U. Montpelier) which binds GalNAc + Glc GIPL in blocking buffer for 72 hrs at 4°C. Membranes were washed with TBS-T 0.1% three times and incubated for one hr at room temperature with goat anti-mouse Ig horseradish peroxidase (HRP)-conjugated antibodies (Southern Biotech, anti-IgM secondary 1:1000 (cat# 1020-05), anti-IgG 1:7500 (cat# 1030-05)). Membranes were then washed with TBS-T 0.1% three times and developed with Immobilon Forte Western HRP Substrate (Millipore, WBLUF0500). All blots were imaged via chemiluminescence on a ChemiDoc Touch (cat#12003153, Bio-Rad). Image Lab 6.1 software (Bio-Rad) was used for analysis of bands.

### NanoLC-MS and MALDI-TOF-MS analysis of GIPLs and GPI-anchored protein glycans released from parasites

Parasite pellets corresponding to approximately 3×10^8^ parasites were harvested as described in 2.2.6 but from 15 T175 flasks and thoroughly rinsed in ice-cold PBS (Corning cat # 21-040 CV), before being pelleted and stored at 80°C. Control, uninfected HFF monolayers were harvested in 1 mM EDTA in PBS and washed 3 times in ice-cold PBS by centrifugation (8 minutes, 2000 x *g*, 4°C). Parasite and host cell pellets were then delipidated, and the protein pellet was extracted with butan-1-ol saturated water to enrich for GPI-anchored proteins, as described (36). GIPL fractions were obtained from the clarified lipid extracts generated during initial pellet delipidation. Fatty acids (FA) were removed from both GPI-anchored proteins and GIPLs by incubation in 0.5 M NH_4_OH (in water for GPI-anchored proteins or after bath sonication in 70% ethanol for GIPLs) under rotation for 6 hrs at 4°C. The glycan core was then released by hydrofluoric acid treatment, C18 cleaned up, re-N-acetylated, permethylated, and dried as described (36).

For nLC-MS, permethylated glycans were redissolved in MeOH and 10μl of were mixed with 90μl 0.1% FA in water. 10μl was injected into a PepMap Acclaim analytical C18 (75 μm, 15 cm, 2 μm pore size) column maintained at 60°C in an Ultimate 3000 RSLC coupled to a Q-Exactive-Plus mass spectrometer (Thermo Fisher Scientific). Column was equilibrated for 10 minutes at 97.5% LC-MS Buffer A (0.1% FA in water) and ramped up to 35% LC-MS Buffer B (80% (v/v) over 2 minutes. Glycan separation was achieved using a linear gradient from 35% to 70% Buffer B over 150 minutes at a flow rate of 300 nl/minutes. The effluent was introduced into the mass spectrometer by nanospray ionization in positive mode via a stainless-steel emitter with spray voltage set to 1.9 kV and capillary temperature set at 275°C. The MS method consisted of a survey Full MS scan at 70,000 resolution in positive ion mode, followed by MS(2) fragmentation of the top 10 most intense peaks using HCD at 40% collision energy and an isolation window of 2 m/z. Dynamic exclusion was set to exclude ions for fragmentation for 30 sec. All the data were processed manually using the Xcalibur 2.0 software package.

For MALDI-TOF-MS, permethylated samples were redissolved in MeOH and 0.5µl aliquots were mixed 0.5µl MeOH saturated with either 2,5-dihydroxybenzoic acid (DHB) or α-cyano-4-hydroxycinnamic acid (CHCA) on the MALDI plate and allowed to air dry. Samples were then analyzed on an ABI 4700 MALDI-TOF-MS operated in positive ion reflectron mode for m/z range 500-5000.

### Flow cytometry of peritoneal exudate cells after infection

Mice were infected i.p. with 10^6^ tachyzoites, and 3 hrs post injection mice were euthanized and peritoneal cavity lavaged with 4mL of 1X PBS and 3 mL of air. The peritoneal exudate fluid was passed through a 70 μm cell strainer, cells were pelleted for staining. Samples were blocked for 30 minutes in FACS buffer (2% FBS 1X PBS) containing Fc Block anti-CD16/32 (2.4G2, BD Biosciences) (1:100 dilution), 5% normal hamster serum, and 5% normal rat serum (Jackson ImmunoResearch). After blocking, cells were stained at a 1:100 dilution for 30 minutes with the following antibodies: CD11b BUV395 (M1/70, BD), Gr.1 PE (RB6-8C5, BioLegend), F4.80 BV421 (BM8, Thermo Fisher), MHC II APC (AF6-120.2, Thermo Fisher), B220 APC-Cy7 (RA3-6B2, BioLegend), CD19 BUV785 (6D5, BioLegend), NK 1.1 PE Cy7 (PK136, Thermo Fisher), CD3ε BV510 (17A2, BD) and Propidium Iodine (PI) (1:1000). After incubation, cells were washed three times in FACS buffer and resuspended in FACS buffer. Samples were run on a ZE5 flow cytometer (Bio Rad) and analyzed using FlowJo software version 10.

### Flow cytometry for parasite antigen expression

Syringe lysed parasites were plated in a micro-titer plate at 8×10^5^ parasites per well and incubated with one of the following antibodies at (1:100) to detect antigen expression: mouse anti-SAG1 (T4 IE5, BEI), mouse anti-SAG3 (T4 1F12, BEI), mouse anti-p35 (T4 3F12, BEI). After 20 minutes incubation on ice, parasites were washed twice with FACS buffer and stained with (1:100) goat anti-mouse IgG-APC (BioLegend, cat# Poly4053). After incubation, cells were washed and resuspended in FACS buffer as described above. Samples were run on a ZE5 flow cytometer and analyzed using FlowJo software version 10.

### Serum antibody parasite binding assay

For serum reactivity analysis, syringe-lysed GFP-expressing strains were fixed in 3% formaldehyde for 20 minutes, washed twice in PBS, and plated in 96-well micro-titer plates at 4×10^5^ parasites/well. The parasites were then incubated with serum from chronically infected mice, in serum concentrations ranging from 10^-2^ to 10^-6^ diluted in FACS buffer, for 20 minutes at 37°C. Parasites were then washed with FACS buffer and placed on ice for incubation with anti-isotype detection antibodies depending on application: anti-IgG3-BV421 (R40-82, BD Bioscience), anti-IgM-PE/Cy7 (RMM-1, BioLegend), anti-IgG1-APC (RMG1-1, BioLegend), anti-IgG2b-PE (RMG2b-1, BioLegend), anti-IgG2a-PerCP/Cy5.5 (RMG2a-62, BioLegend). Parasites were washed and resuspended in FACS buffer as described before. Flow cytometry was performed on an LSR analyzer (BD) and analyzed using FlowJo software version 10. The MFI was computed by gating on forward and side scatter characteristics consistent with parasites and further gated for GFP+ if the parasite expressed GFP using FlowJo.

### Complement C3b binding assay

For analysis of C3b binding, parasites were syringe lysed, washed, and resuspended to a concertation of 1×10^7^ parasites per mL in Hank’s Balancing Salt Solution Buffer (HBSS) (ThermoFisher, cat#14175095). Parasites were plated in a 96-well plate at 10^6^ parasites per well with 10% blood plasma from naive mice and incubated at 37°C for 30 minutes. After incubation, complement activation was stopped by washing with cold 1X PBS. Parasites were spun down at 611 x g for 3 minutes and washed 3 times using cold 1X PBS. Parasites were then fixed using 3% formaldehyde for 10 minutes and then washed using cold 1X PBS. Parasites were then stained with mouse anti-C3b (10C7) (Invitrogen MA1-70054) (1:200) for 20 minutes. After primary stain, parasites were washed with cold 1X PBS and stained with anti-mouse IgG1 APC (RMG1-1, BioLegend) (1:100) for 20 minutes on ice. After incubation, parasites were washed 3 times and resuspended in 1X PBS for analysis. Samples were run on a ZE5 machine and data analyzed using FlowJo version 10.3. MFI was determined as described above.

### Galectin-3 binding assay

Recombinant Gal-3 (R&D Systems, Cat# 1197-GA-050) was pre-diluted in phosphate-buffered saline (PBS) containing 10% bovine serum albumin (BSA) and 14 mM β-mercaptoethanol. Parasites were syringe lysed, washed, and 8×10^6^ parasites were incubated with either 10 or 5 µg of recombinant Gal-3 for one hour at 4°C in FACS buffer. Parasites were washed to remove unbound Gal-3 and incubated with a PE-conjugated anti-Gal-3 antibody (eBioM3/38 (M3/38), Invitrogen) at a 1:100 dilution in FACS buffer for 20 minutes at 4°C. After two further washes, the samples were analyzed using a ZE5 Cell Analyzer (Bio-Rad) and data processed as described above.

### CD36-Fc binding assay

10^7^ parasites were incubated with 0.5 μg recombinant human IgG Fc (R&D Systems, cat# 110-HG) or recombinant CD36-Fc (R&D Systems, #2519-CD) in 50μl binding buffer (0.14 M NaCl, 2.5mM CaCl_2_, 0.01M HEPES (pH7.4), 3% BSA) for 1 hr at 15°C. Parasites were washed once to remove unbound recombinant protein and the parasites were then incubated with mouse anti-SAG1-AF405 (TP3cc, Novus Biologicals) and donkey anti-human IgG-Daylight 550 (Thermo Fisher, SA5-10127). The stained parasites were acquired by Attune NxT flow cytometry. The CD36-Fc MFI was computed by gating on GFP+/SAG1+ parasites using FlowJo.

### Opsonization and invasion assays

Bone marrow-derived macrophages (BMDMs) were generated in L929 conditioned medium from female C57BL/6J mice as previously described (37). BMDMs were plated a day in advance on a coverslip lined 24-well plate at 8 x10^5^ cells in 1mL per well, in “BMDM media” supplemented with 10% L929 (20% HI FBS, 1% penicillin-streptomycin, 1% non-essential amino acids (ThermoFisher, cat#11-140-050), 1% sodium pyruvate). The day of the experiment, freshly lysed parasites were washed and resuspended for an MOI of 0.5 in Toxo media warmed to 37°C. In a 96-well plate, serum dilutions were made in Toxo media that was pre-warmed to 37°C. Parasites were added to the diluted serum and incubated at 37°C for 20 minutes before being added to the BMDM coverslip wells with fresh Toxo medium. The BMDM plates were spun at 304 x g for 5 minutes to synchronize parasite invasion between wells and incubated at 37°C for 40 minutes to allow for phagocytosis or invasion. After 40 minutes, media was replaced with 300μl of 3% formaldehyde in 1X PBS, fixed for 20 minutes, washed three times with 1X PBS, blocked and permeabilized with “Blocking Buffer” for 1 hr (1X PBS with 10% normal goat serum, and 0.01% saponin). Cells were stained with rabbit anti-GRA7 (1:5000) (gift from John Boothroyd, Stanford) and rat anti-LAMP1 (1D4B, BD Biosciences) (1:500) diluted in blocking buffer for 1 hr or overnight at 4°C. After primary antibody incubation, wells were washed three times with 1X PBS before secondary antibody incubation. Secondary antibodies were goat anti-rabbit AF647 (1:3000) (Thermo Fisher, cat#A21245) goat anti-rat AF594 (1:1000) (Thermo Fisher cat#A11007), and DAPI (1:10000), and if parasites were not GFP-expressing, goat anti-Toxo conjugated FITC (1:400) (ViroStat, cat#0283) was also used. Fluorescence microscopy was performed (Nikon Eclipse T*i*-5) and images were captured at 60x and blinded for analysis. Images were processed (Nikon Elements) and 80-100 GFP+ events were quantified for parasite association with Lamp1+ or for containment within a ring-like GRA7+ parasitophorous vacuole (PV); fraction opsonization was calculated as (Lamp1+GFP+ counts / Lamp1+GFP+ plus GRA7 PV+GFP+ total counts).

For invasion assays BMDMs were plated on coverslips in 24-well plates overnight as described above and infected the next day at an MOI of 0.25 parasites. Plates were spun at 304 x g to synchronize invasion and incubated at 37°C for 20 minutes before washing once in 1X PBS and fixing in 3% formaldehyde. Cells were not permeabilized. Wells were washed and stained with mouse anti-SAG1 (1:5000) for 1 hr. Cells were washed, and secondary antibodies goat anti-mouse AF594 (1:3000) and DAPI (1:1000) were used for 1 hr. Fluorescence microscopy was performed as described above, with 100 GFP+ parasites counted per slide and quantified for parasite for SAG1+ staining. Invaded parasites were determined to be GFP+ SAG1-. Fraction of invasion was calculated as (SAG1-GFP+ counts / total GFP+ counts).

### Neutralization assay

MEFs were plated a day in advance in a 96-well plate at 5×10^5^ cells per well in MEF media. The day of the experiment, freshly lysed parasites were washed and resuspended to a concentration of 1.6×10^7^ in in Toxo media warmed to 37°C. In a 96 well plate, blood plasma 1:10 dilution was made in Toxo media pre-warmed to 37°C and incubated at 37°C in 10% serum for 20 minutes before being added to the MEFs, such that in the absence of serum the MOI is approximately 0.5. The plate was spun at 304 x g for 3 minutes to synchronize parasite invasion and incubated at 37°C for 2 hrs. Supernatant was carefully removed, and 20μl of trypsin was added to each well and incubated at 37°C for 5 minutes to dislodge cells from the plate. Cells were harvested with 100μl of cold 1X PBS, transferred to a FACS plate and washed 3 times with FACS buffer to remove trypsin. Cells were resuspended in 1:1000 PI in FACS buffer, and then analyzed via flow cytometry for GFP+ PI-MEFs to indicate invasion. Neutralization was defined as a ratio: the percentage of PI-cells infected with parasites incubated with serum divided by the percentage of PI-cells infected with parasites without serum incubation. For non-GFP+ expressing parasites, cells were fixed after trypsinization in 3% formaldehyde, washed, and blocked for 1 hr before staining with anti-Toxo-FITC (ViroStat, cat#0283). Neutralization ratios were determined as described, but PI was not included in the discrimination. Samples were run on a ZE5 machine and data analyzed using FlowJo software version 10.

### Parasite lytic cycle

For plaque sizes, HFFs were plated in 24-well plates and allowed to grow to confluency before adding 100 or 300 syringe-lysed parasites and allowed to grow for 5 days. Plaque sizes were measured (Nikon Elements) under 4x objective (Nikon Eclipse T*i*-5).

For parasites per vacuole assays, 8×10^5^ MEFs were plated on coverslips in 24-well plates and allowed to adhere before infecting with 10^5^ parasites. Plates were spun at 475 x g to synchronize invasion and incubated for 15 minutes at 37°C. After 15 minutes cells were washed twice to remove any unattached parasites. Fresh Toxo media was added, and plates incubated at 37°C for 16 hours before being fixed, blocked with 0.1% saponin blocking buffer (as described above), and stained for 1 hr with rabbit anti-GRA7 (1:3000). Cells were washed and incubated with secondary goat anti-rabbit AF 594 (1:3000) (Thermo Fisher, cat#A11037) and DAPI (1:10,000) for 1 hr. 100 GFP+ vacuoles per slide were quantified for number of parasites residing in GRA7+ vacuoles under 100X objective (Nikon Eclipse T*i*-5).

For attachment time course assays, HFFs were plated in 24-well plates and allowed to grow to confluency. 200 syringe lysed parasites were added per well and plates were spun at 475 x g to synchronize invasion and incubated for 30, 60, or 90 minutes before washing 5 times in 1X PBS to remove any unattached parasites. Fresh Toxo media was added and plates were incubated at 37°C for 5 days before counting plaques under the microscope. Plaque numbers were normalized to those obtained from wells that did not undergo any washes.

### Quantitative PCR for cytokine expression and parasite burden

Cells were collected from mouse peritoneal lavages as described above and spleens. Cells were filtered through a 70 μm filter and centrifuged 611 x g for 7 minutes to pellet and washed with sterile 1X PBS before pelleting again. Samples were removed of red blood cells through treatment of ACK lysis buffer (0.5 M NH_4_Cl, 0.01 M KHCO_3_, .0001 M EDTA) for 5 minutes before being washed with 1X PBS. Cells were stored in RNA-later (ThermoFisher, cat#AM7021) in -80°C until RNA isolation. After thawing samples, aliquots were removed and diluted with 1X PBS and centrifuged to pellet cells and remove RNA-later. RNA was isolated using the RNEasy Mini Kit (QIAGEN, cat#74134) and cDNA was synthesized using the Invitrogen High-Capacity cDNA Reverse Transcription Kit (ThermoFisher, cat#4368814). Quantitative PCR was performed on synthesized cDNA using iTaq Universal SYBR Green Supermix (BioRad, #172511). Normalization of all samples was calculated in comparison to *Actb* expression levels. Fold change in cytokines were determined through the 2^-ΔΔCt^ method.

For parasite burden determination by quantitative PCR, peritoneal exudate and spleen cells, liver and lung tissues were harvested, and appropriate amounts were digested according to manufacturer’s protocols using the QIAGEN DNeasy Blood & Tissue Kit (#69504). Brains were isolated and DNA purified as follows. In brief, whole brains were harvested in 5 mL of 1X PBS and homogenized through an 18G needle of a 10mL syringe and passed up and down through the needle 10 times before being spun down at 611 x g for 7 minutes to pellet. 100 μL of brain homogenates were processed using the QIAGEN DNeasy Blood & Tissue kit according to the manufacturer’s directions. Standard curves were made from known numbers of isolated parasite DNA ranging from 10^1^ to 10^6^ parasites per well. Isolated DNA samples were amplified targeting the *Toxoplasma-*specific B1 gene, and the standard curve was used to quantify parasite number in each sample per μg of isolated DNA.

### Statistics

Statistical analyses were performed with GraphPad Prism 8 software. Statistical significance was defined as P < 0.05. P values between two groups were calculated using paired or non-paired two-tailed t-tests. One or two-way ANOVA was used for comparison across more than two groups with Tukey or Dunnett correction, as recommended by Prism. Survival curve significance was calculated using log-rank Mantel-Cox testing.

## RESULTS

### *Tg_207750* and *Tg_266320* are predicted to be the *T. gondii* GPI sidechain-modifying glycosyltransferases, PIGJ and PIGE, respectively

*T. gondii* strains are known to differ in virulence in various intermediate hosts (38). In laboratory mice, type III strains are referred to as avirulent because they require a dose of over 10^5^ parasites to achieve 50% lethality (38,39). Meanwhile, type I strains are highly virulent, and one parasite is enough to be 100% lethal in naïve mice (40,41). However, when C57BL/6J mice given a primary infection with the avirulent type III strain CEP and allowed to progress to a chronic infection, they survive secondary infections (i.e., “challenge”) with high dose type I strain RH parasites due to protective immunological memory responses that are generated following the initial parasite exposure (42–44). In contrast, when mice infected with CEP are challenged with the type I strain GT1 they succumb (43). RH and GT1 are highly similar type I strains and the gene *Tg_207750* was noted to be six times more highly expressed in RH compared to GT1 (45). Based on sequence analysis, *Tg_207750* was predicted to be a glycosyltransferase that adds an acetylated hexose (HexNAc) to a core mannose of complex N-glycans (45). Moreover, *Tg_207750* was one of two *T. gondii* βHexNAc transferase-like GTs we proposed to be responsible for adding the GalNAc to the GPI of *T. gondii* (36). Thus, we hypothesize that *Tg_207750* is partially responsible for differences in secondary infection virulence between RH and GT1 type I strains and the GPI GalNAc transferase we name here as “PIGJ” (**Fig 1A**).

**Figure 1:**
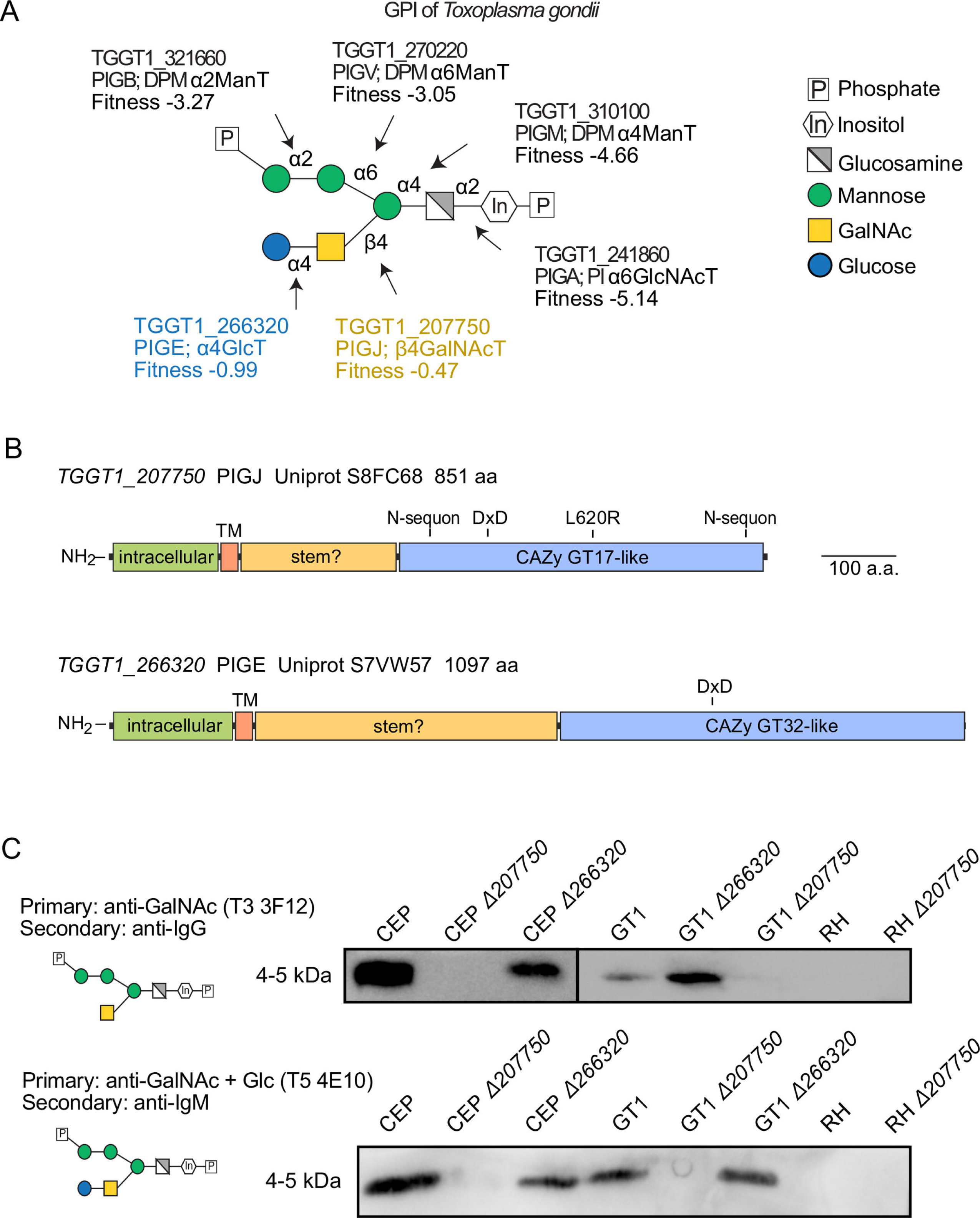
Glycoform specific antibodies implicate *Tg_207750* as the GalNAc GPI sidechain modifying enzyme, PIGJ. A) Schematic of the T. gondii GPI with known and putative glycosyltransferases involved in the GPI synthetic pathway. Transferases show their gene ID, gene name, transferase type, fitness score (the more negative the fitness score being more lethal to the parasite when inactivated) (4), and the linkage they form. The two novel glycosyltransferases identified and characterized in this study are labeled in color and named ‘PIGJ’ and ‘PIGE’ responsible for the GalNAc and Glc additions, respectively. B) Gene models for *Tg_207750* (PIGJ) and *Tg_266320* (PIGE) from ToxoDB were translated and subjected to BLASTp and other homology searches to predict the evolution and function of the proteins. Regions of predicted function are represented: TM transmembrane, putative stem region, DxD catalytic site, N-sequon potential glycosylation sites and a non-synonymous SNP at position 620 in PIGJ between GT1 (L) and RH (R) type I strains. C) Representative western blots confirming the loss of the sidechain in *Δ207750* mutants through sidechain specific antibody reactivity to the low molecular weight (<10 kDa) GIPL antigen (Clones: T3 3F12 binds to the GalNAc glycoform and T5 4E10 prefers the GalNAc + Glc glycoform of GIPL). GIPL from CEP and GT1 *Δ266320* mutants are recognized by both glycoform-specific antibodies. GIPL from RH was not detected by T3 3F12 nor T5 4E10. Representative of 3-5 experiments.

Informatic analysis revealed *Tg_207750* is predicted to be a type 2 transmembrane protein (46) with a C-terminal glycosyltransferase domain with greatest similarity to CAZy family GT17 (36) (**Fig 1B**). The occurrence of a non-synonymous SNP within the GT17 CAZy-like domain was also noted between RH and GT1. Reciprocal BLASTp studies indicate that PIGJ is coccidian specific and not found in *Plasmodium* or *Cryptosporidium spp.*, or Piroplasm apicomplexans (not shown). The catalytic DxD domain is separated from the transmembrane domain by a stretch of about 170 aa, which in other GTs comprises a stem-like region that may mediate protein interactions or appropriately orient the catalytic domain. The 143-aa cytoplasmic region is predicted to be disordered based on amino acid composition. AlphaFold predictions confirm the potential of the GT17 region to fold as a GT-A superfamily domain, but do not confidently predict a structure for the stem-like or cytoplasmic regions. Now referred to as PIGJ, *Tg_207750* evolved independently of PGAP4, the GT that catalyzes the same reaction in animals (30). PGAP4 is a CAZy GT109 family member that bears no recognizable homology with CAZy GT17 sequences and has a distinct architecture that includes a transmembrane hairpin inserted within the canonical catalytic domain. Thus, formation of the same GalNAcb1,4Man linkage in coccidian apicomplexans and in animal hosts is the result of convergent evolution.

The gene *Tg_266320* was previously predicted to encode the GPI αGlcT that mediates addition of the terminal Glc to GalNAc to complete the disaccharide sidechain (36) (**Fig 1A**). *Tg_266320* is also predicted to be a type 2 transmembrane protein (**Fig. 1B**). Now referred to as “PIGE”, *Tg_266320* has greatest similarity to CAZy GT32 glycosyltransferases (36). BLASTp searches suggest that this protein is likewise restricted to coccidian apicomplexans (not shown). The most similar GT found is lactosylceramide 4-alpha-galactosyltransferase (a.k.a. globotriaosylceramide or Gb3 synthase), which attaches αGal to the 4-position of a β-linked Gal acceptor across many eukaryotes (47). Thus, the linkage formed is the same, but the donor and acceptor sugars are distinct, which is an evolutionary variation common among CAZy GT32 GTs. GT32 GTs utilize sugar nucleotide rather than Dol-P-sugar donors, as previously documented for the enzyme that assembles the Glcα1,4-linkage in the *Toxoplasma* GPI (48). Though PIGE and Gb3 synthase are both type 2 membrane proteins that modify glycolipids, PIGE is distinctive with its ∼385 aa long stem-like region and a ∼160 aa poorly conserved, likely disordered cytoplasmic region, whereas Gb3 synthase is only 360 aa overall with a negligible cytoplasmic region. Given evidence for multiprotein complexes in the GPI biosynthetic pathways of other organisms (49), the cytoplasmic and stem-like regions of PIGJ and PIGE might integrate them into complexes for orderly and efficient sequential GPI-assembly in the rER (50). This hypothesis is consistent with the lack of KDEL-like rER retentions signals at their C-termini.

### Western blot analysis implicates *Tg_207750* as PIGJ

To address their roles in the GPI sidechain synthetic pathway, each predicted GT gene was targeted for disruption by CRISPR-Cas9 and repair with a drug selectable marker (**Fig S2 S3**). Mutants of each gene were made in both type I (RH, GT1) and type III (CEP) genetic backgrounds (**Table S1**), and integration of the selectable markers was confirmed with diagnostic primers and PCR (**Fig S2 S3**). Using glycoform-specific antibodies that bind the protein-free GPI lipid (GIPL) sidechains (17), western blot analysis revealed these antibodies lose recognition of both sidechain glycoforms derived from the GT1 *Δ207750* and CEP *Δ207750* strains (**Fig 1C**). This indicates that the enzyme encoded by *Tg_207750* is likely responsible for GalNAc addition to the mannose core of the GPI. Western blot analysis of GT1 *Δ266320* and CEP *Δ266320* mutants retained GIPL recognition by the T3 3F12 GalNAc GIPL glycoform specific antibody, and in the case of GT1 *Δ266320* increased detection was routinely observed. In contrast, recognition of the GalNAc + Glc glycoform by the T5 4E10 antibody was unimpeded against GIPL from GT1 *Δ266320* and CEP *Δ266320* mutants (**Fig 1C**). A technical note of importance, while the T5 4E10 antibody has preferential reactivity to GalNAc + Glc compared to GalNAc GIPLs of *T. gondii* (17), it can detect GalNAc sidechain bearing GIPLs of human intermediate GPIs (29), suggesting significant cross-reactivity to antigen by this antibody clone.

To further explore the recognition potential by these antibodies, we decided to monitor their ability to bind intact parasites by flow cytometry. Like that of our western blot analysis, T3 3F12 and T5 4E10 staining of the GT1 *Δ207750* mutant was significantly reduced compared to the parental parasite strain. However, the staining was not entirely abrogated (**Fig S4**). Regarding the GT1 *Δ266320* strain, T3 3F12 recognition of the mutant parasite is enhanced suggesting increased presence of the GalNAc bearing GPI sidechains. In contrast, T5 4E10 recognition of GT1 *Δ266320* is only marginally reduced, calling into question whether this GT is responsible for adding the terminal Glc to the GPI sidechain, at least using this antibody clone which has significant cross-reactivity to GPI antigens (17,29). Finally, while *Tg_207750* is more highly expressed in RH compared to GT1 type I strains (45,51), we do not detect GIPL in the RH strain via western blot using these antibodies (**Fig 1C**), nor do we detect the disaccharide sidechain by mass spectrometry of GPI anchors isolated from this strain (36). Consistent with these observations, reduced T5 4E10 and T3 3F12 binding of the RH strain relative to the GT1 strain is observed by flow cytometry, with only slight reductions in T3 3F12 staining of the RH *Δ207750* compared to the parental RH strain is seen (**Fig S4**).

### Structural analysis confirms GPI sidechain glycosyltransferases of *T. gondii*

Given the low detection of GIPL in the RH background by GIPL-specific antibody and mass spec, the GT1 strain was subjected to GPI sidechain structural analysis via mass spectrometry (**Fig 2**). A previous study developed a new method to isolate and analyze the glycan component of GPI anchors using mass spectrometry (36). The method first enriched for GPI-anchored proteins using butan-1-ol extraction of material delipidated by extraction with CHCl_3_ and MeOH. The fraction was saponified to remove potential fatty acids (FAs), and the glycan core was released by phosphodiester bond cleavage with hydrofluoric acid (HF). Finally, the glycan was N-acetylated and permethylated, and analyzed initially by MALDI-TOF-MS. The most abundant ion from the parental GT1 strain corresponded to a glycan with the composition of four hexose residues, two HexNAcs, and one hexitol, or H4N2Ino (**Fig 2A**). This composition corresponds to the previously described structure consisting of three Man, one Glc, one GalNAc, and one inositol, as shown in Figure 1A. In addition, less abundant ions were found that correspond to H3N2Ino and H3N1Ino, suggesting the presence of isoforms with a monosaccharide side arm, or none at all. Ions whose composition corresponded to multimers of hexose, H3, H4 and H5, were also detected.

**Figure 2:**
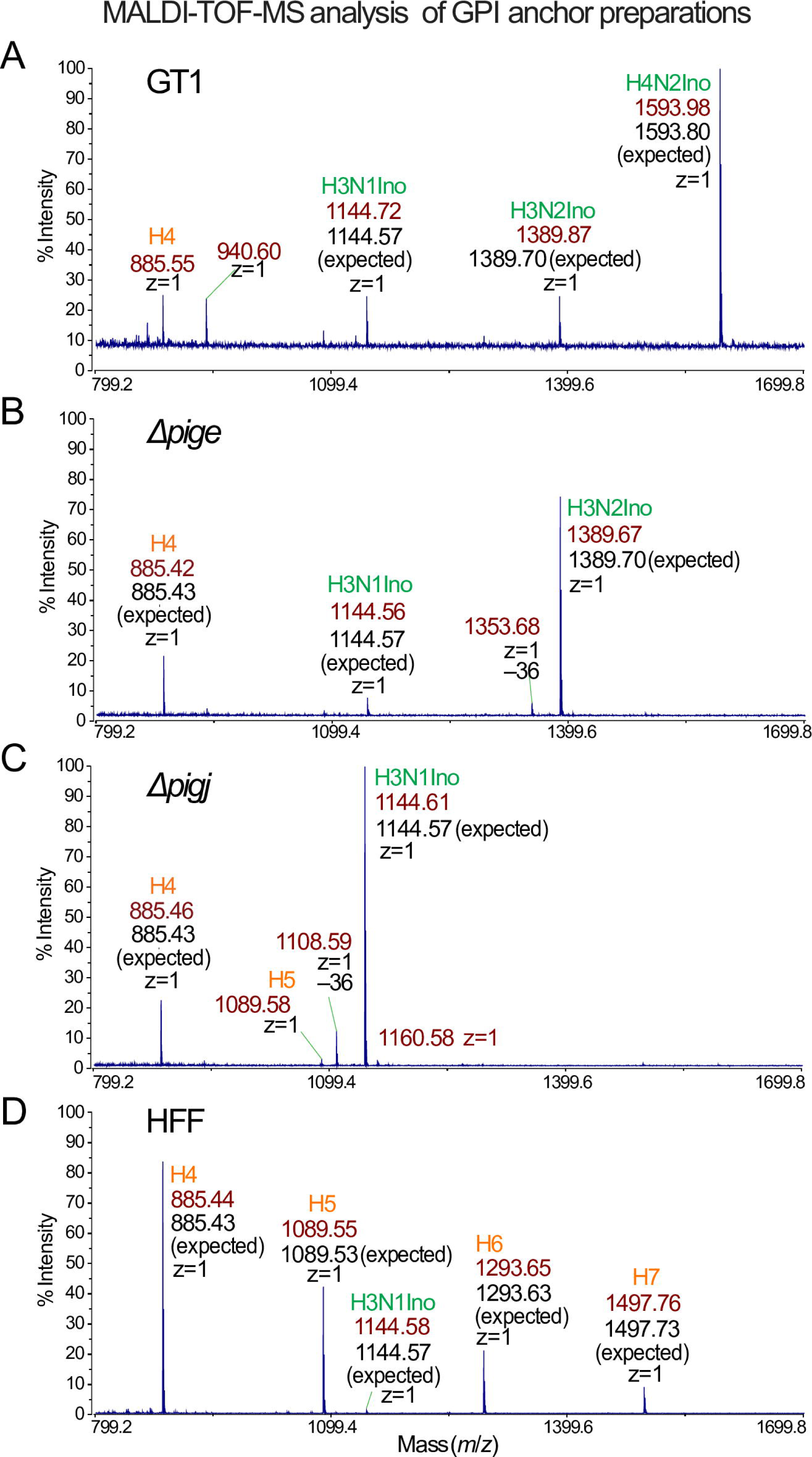
MALDI-TOF-MS analysis of GPI preparations from GPI-anchored proteins reveal the identities of PIGE and PIGJ as GPI sidechain glycosylransferases. Glycans isolated from GPI-anchored proteins from tachyzoite stage parasites were N-acetylated, permethylated, mixed with CHCA matrix and analyzed in reflectron and positive ion mode. The m/z range of 800-1700 is shown. Ions that correspond to singly charged (z=1) sodiated GPI glycans are labeled in green, with the observed monoisotopic m/z value (in dark red), and the predicted m/z values for the assignment (in black). As an example, the composition assignment for an ion possessing three hexoses (known to be mannoses), one HexNAc (known to be N-acetylglucosamine), and one hexitol (known to be inositol) is represented as H3N1Ino. Ions that correspond to Hex(n) oligomers are labeled in red. A) Parental GT1 strain. B) *Δpige*. C) *Δpigj*. D) Host cells (HFFs).

Since parasites are grown in HFFs, we assessed the potential contribution of HFFs to these ions, since mammals also express GPI anchor glycans with a disaccharide side chain on the α4-linked (first) Man residue (29). Preliminary analysis of N-glycans, performed as described in Gas-Pascual et al. (2019), showed the presence of N-glycans from HFFs as well as tachyzoites (data not shown), such that >25% of the sample was potentially of HFF origin. Therefore, a GPI-anchor fraction from HFFs was examined. As shown in Figure 2D, this sample contained a series of hexose oligomers whose abundance decreases with length, which potentially represent breakdown products of glycogen, as well as an ion corresponding to H3N1Ino. Previous analysis of a preparation of spontaneously lysed extracellular tachyzoites that were unlikely to be contaminated by host cells, and that were washed extensively to minimize serum contaminants, yielded no detectable hexosamers (36), consistent with the low level of starch accumulation at the tachyzoite stage. However, the ratio of H3N1Ino to Hex4 was small in HFFs compared to the large ratio in the GT1 sample (**Fig 2A, D**). If the amount of H4 is used as a measure of contamination of HFFs to the parasite to sample, it is evident that the great majority of the GPI glycans originated from the parasites.

To verify these interpretations, the GT1 GPI preparation was also examined by MS analysis in an Orbitrap mass spectrometer after separation by C18 nano-HPLC. As shown in Figure 3, the elution profile of all ions (base peak chromatogram) from the GT1 strain included a prominent peak eluting at 49.04 min (**Fig 3A**). This peak coincided with the extracted ion chromatogram analysis (EIC) of H4N2Ino in the lower trace of panel A. A second peak eluting at about 33.77 min corresponded to the elution positions of H3N1Ino and H3N2Ino, which were not well resolved by this method. These ions eluted as a mixture of primarily singly charged H^+^ or NH_4_^+^ ions, and doubly charged ions in either 2H^+^ or H^+^/NH_4_^+^ states, as shown in the right-hand panels. These ions matched the expected *m*/*z* values with 1-2 orders of magnitude higher accuracy compared to the MALDI-TOF-MS method. Furthermore, collision-based decomposition (HCD MS-2) revealed a series of daughter ions that confirmed that it consisted of various combinations of monosaccharides including non-reducing end Hex residues (**Fig S5A**) that are completely consistent with structure shown in Figure 1A. As expected, all 3 GPI isoforms were detected, but at a ratio of 1.0:0.25:0.13 for H4N2Ino:H3N2Ino:H3N1Ino, respectively (**Fig 3A**), consistent with the MALDI-TOF-MS data (**Fig 2A**). Most of the remaining ions in the base peak chromatogram matched the elution positions of the H3, H4 and H5 hexosamers, as described for the HFF GPI sample (**Fig 3D**). Note that these isomers elute in pairs with the same *m*/*z* value, indicating that they represent a mixture of reducing end α- and β-anomers with distinct elution times. While all three GPI isoforms were also detected in the HFF samples, they were of very low relative abundance and H3N1Ino rather than H4N2Ino was the most abundant ion. Taken altogether, the results confirm the presence of the previously described Glc-GalNAc-side chain of parasite GPI anchors, and good evidence for isomers with only GalNAc or no side chain at all.

**Figure 3:**
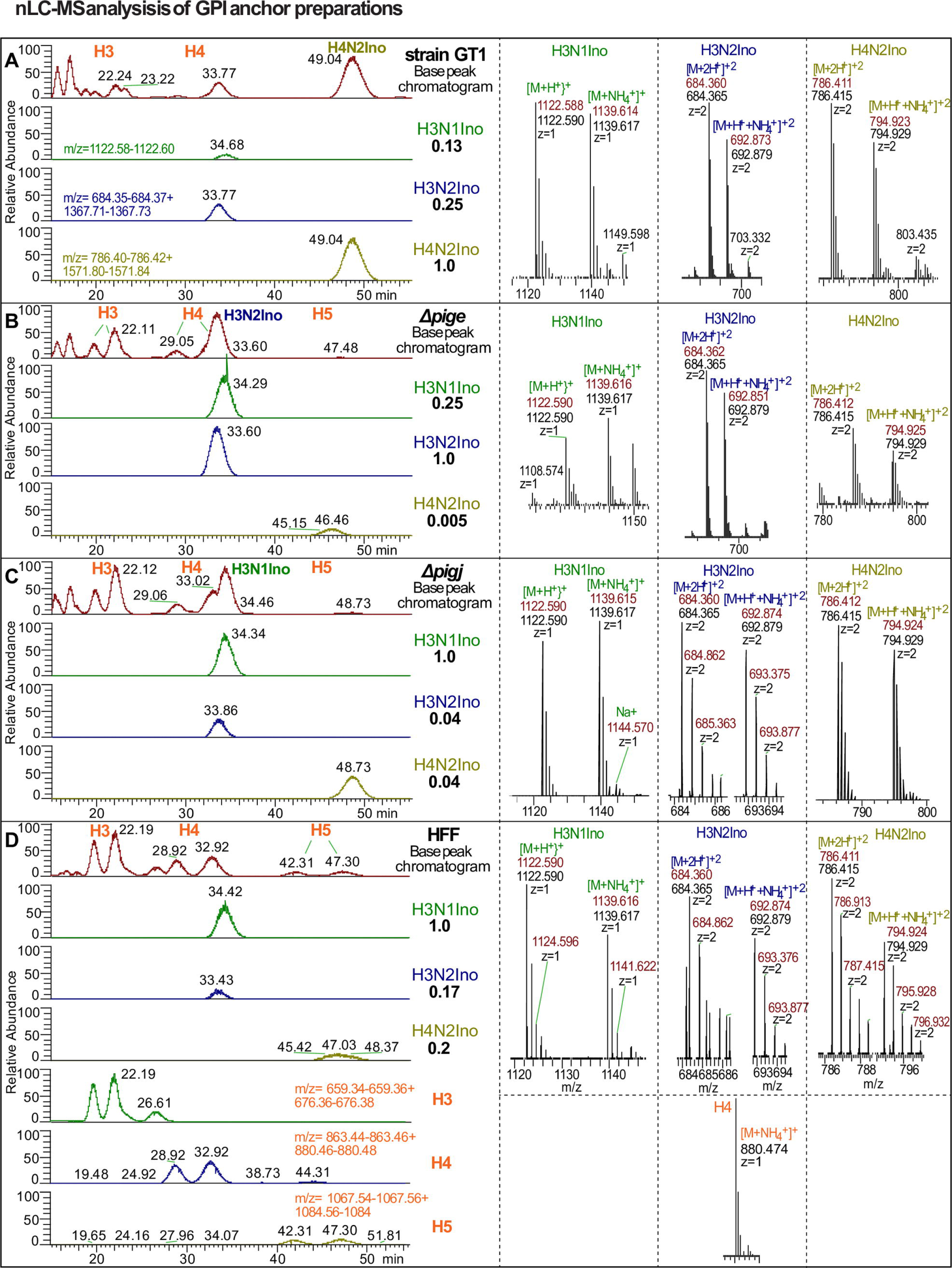
nLC-MS analysis of GPI-anchor preparations confirm the GPI sidechain transferase activity of PIGJ and PIGE in *T. gondii*. Isolated GPI permethylated glycans described in Figure 2 were reanalyzed by nLC separation on a C18 column and hyphenated analysis in an Orbitrap mass spectrometer in positive ion mode. The left column of panels shows base peak chromatograms (all ions m/z 500-2000) and extracted ion chromatograms (EIC) for each of the indicated targets (m/z ranges are specified in panels A and D). The ratio of ion intensities over the EIC m/z ranges shown for H3N1Ino, H3N2Ino, and H4N2Ino are shown relative to the most abundant ion. The most abundant ions are labeled in the base peak chromatogram. The right column of panels shows representative mass spectra of the two most abundant ions eluting with the target and used for quantitating relative levels. Levels of hexosamers were not quantitated. Observed m/z values are in dark red, expected values are in black, and the charge state is as indicated. A) Parental GT1 strain. B) *Δpige*. C) *Δpigj*. D) Host cells (HFFs).

To confirm the predicted role of PIGE as the GPI Glc transferase, the GPI fraction from *Δpige* parasites was analyzed as above. Strikingly, only the H3N2Ino and H3N1Ino isoforms were detected, along with substantial levels of H4 and H5 (**Fig 2B**). Using the more sensitive nLC-MS method, the full-length H4N2Ino isoform was detected but at 0.005% of the level of H3N2Ino (**Fig 3B**), which is likely a contaminant from the HFF cells. To confirm the predicted role of PIGJ as the GPI GalNAc transferase, the GPI fraction from *Δpigj* parasites was similarly analyzed. In this case, essentially only the H3N1Ino glycan was detected together with substantial H3 and H4 glycans (**Fig 3C**). The trace levels of H4N2Ino and H3N2Ino glycans detected were readily explained as carryover from the host HFF cells as indicated by the abundant H3 and H4 glycans. Both mutant forms of the GPI-anchor fragmented to yield expected glycan products (**Fig S5B, C**).

*T. gondii* GIPLs have been described to possess the same Glc-GalNAc-sidearm on the same α4-linked core Man residue. To confirm the findings described above for GPI-anchors, the method for isolating GPI-glycans was applied to the organic fraction obtained from the initial delipidation step used to isolate GPI-anchored proteins, as described in Material and Methods. Using the same MALDI-TOF-MS method as above though with a different matrix, a series of ions was observed that suggested the presence of H4N2Ino, H3N2Ino, H3N1Ino, and H4 (**Fig S6A**). However, the m/z values for the parasite glycans were 36 units smaller than expected, whereas H4 had the expected value. In comparison, a parallel analysis using a different matrix yielded the expected m/z for the most abundant ion (**Fig S6A inset**) albeit also substantial -36 ion. In addition, ions at -14 units were detected at low abundance, suggesting slightly incomplete methylation. To examine whether these mass defects were an artifact of MALDI-ionization, the sample was also analyzed by nLC-MS. Significantly, only ions corresponding to the expected values were detected (**Fig S7A**), and ions with mass defects of -36 and -14 were insignificant. Therefore, these defects, which were also present at trace levels in the GPI-anchor analysis (**Fig 2B, C**) but not originally noted, were evidently artifacts of the MALDI ionization method. The ratio of H4N2Ino:H3N2INo:H3N1Ino (1.0:0.33:0.17) was like that of the GPI isoform ratios observed in the MALDI-TOF-MS method.

Analysis of the GIPL fractions from *Δpige* and *Δpigj* strains yielded the same findings as from the GPI-anchor studies. Using either MALDI-TOF-MS (**Fig S6B**) or nLC-MS (**Fig S7B**), the *Δpige* sample contained primarily the monosaccharide side chain with negligible disaccharide side chain presence. Similarly, the *Δpigj* sample lacked the side chain altogether (**Figs S6C, S7C**) with the miniscule levels of side chain isoforms attributable to host cell contamination (**Figs S6D, S7C**). Thus, the same glycosyltransferases act to generate the Glc-GalNAc-side-arm in both GPI-APs and GIPLs, and a significant fraction of glycans contain a monosaccharide or no side-arm in the wildtype GT1 strain as well. In summary, mass spec results confirm the predicted role of both glycosyltransferases. *Δ266320*, the gene hereon referred to as PIGE, lacks the GalNAc + Glc glycoform, and *Δ207750*, the gene hereon referred to as PIGJ, completely lacks both glycoforms of the sidechain (GalNAc, GalNAc + Glc) in the GT1 background.

### PIGJ mutants have increased primary and secondary infection virulence

With the knowledge that PIGJ and PIGE are the *T. gondii* GPI sidechain GTs, the opportunity presented itself to test the role of distinct GPI glycoforms in microbial virulence. To this end both primary and secondary infections were performed using the PIGJ and PIGE mutant strains. Importantly, primary infections with the CEP *Δpigj* strain resulted in 60% mortality compared to 0% mortality following wildtype infections (**Fig 4A**). In contrast, mice infected with CEP *Δpige* survived primary infections (**Fig 4A**). Secondary infections were performed in mice chronically infected with the type III strain, and as previously reported (43,44), these mice survive type I RH challenge due to the formation of protective immunity (**Fig 4B**). Importantly, RH *Δpigj* but not RH *Δpige* mutants caused lethal secondary infections. Chronically infected C57BL/6J mice were also challenged with type I GT1 mutants and succumbed similarly to all three strains: GT1, GT1 *Δpigj,* and GT1 *Δpige* (**Fig S8**). Given C57BL6/J mice are susceptible to GT1 challenges (43), it is difficult to observe virulence increases in this context. To explore this further, the virulence phenotypes of the GPI sidechain mutants were screened in A/J mice, which are genetically resistant to primary infections with low virulence type II strains (52) and secondary type I GT1 infections (53). As observed in C57BL/6 mice, CEP *Δpigj* caused lethality during primary infections in A/J mice (**Fig 4C**). During secondary infections with the RH and GT1 GPI sidechain mutants, however, A/J mice were still resistant (**Fig 4D**). Hence, the *Δpigj* deletion produces a virulence phenotype that overcomes the genetic basis for resistance to primary but not secondary infections in A/J mice. Collectively, these data suggest that the GPI sidechain of *T. gondii* prevents lethal outcomes in its hosts, and that the virulence phenotype is less impacted by the terminal Glc residue transferred by PIGE. Furthermore, the data is consistent with the original hypothesis that differential *Tg_207750* gene expression contributes to the type I strain differences in secondary infection virulence, at least in the susceptible C57BL/6 genetic background.

**Figure 4:**
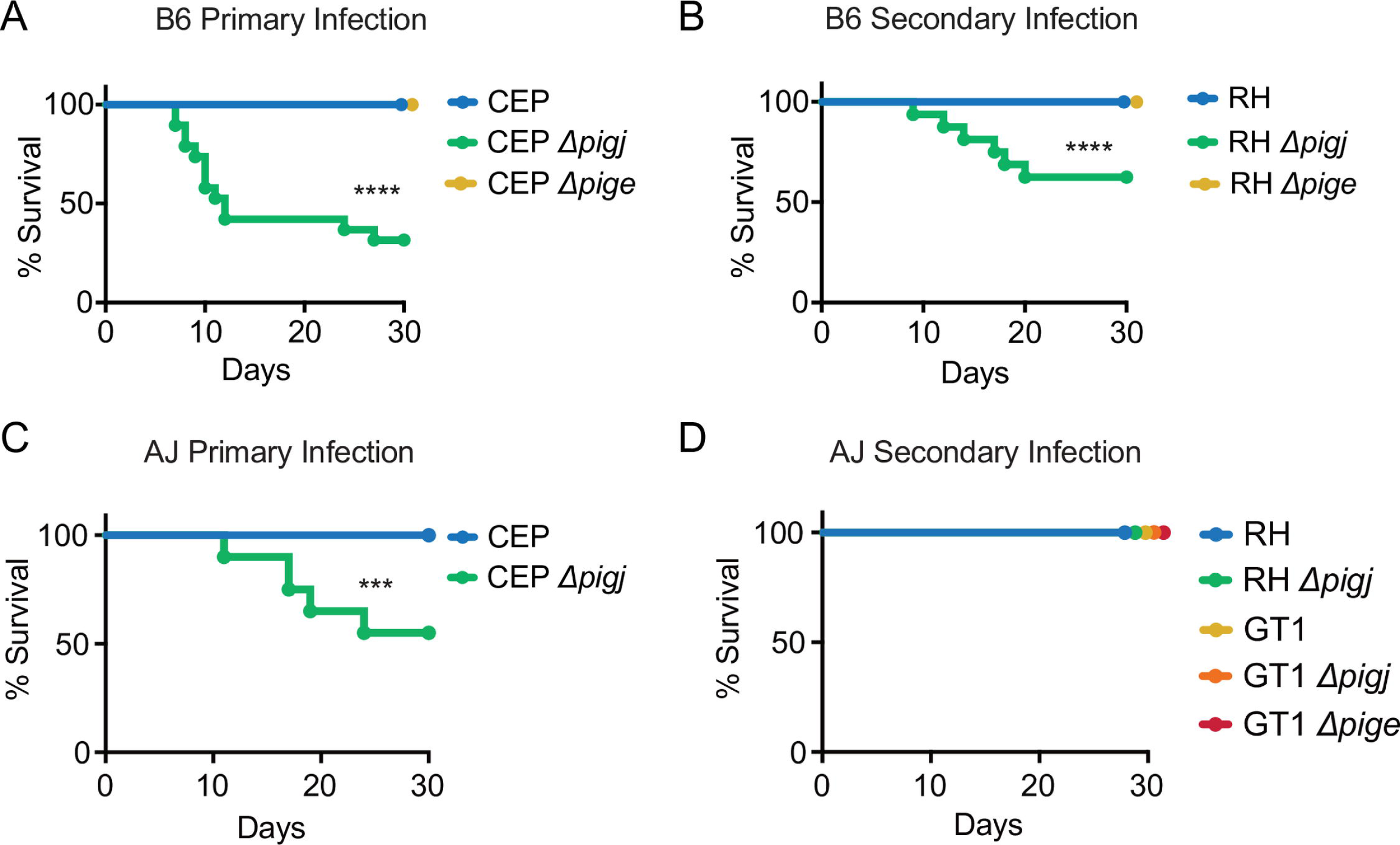
Deletion of the GPI sidechain glycosyltransferase PIGJ but not PIGE results in increased parasite virulence. A) C57BL/6J (B6) mice were given primary infections of 10^4^ parasites (i.p.) of the indicated variants of the avirulent type III CEP *hxgprt-GFP:cLuc* strains. Cumulative survival from 2-4 experiments is plotted (mice; n = 20 CEP, 19 CEP *Δpigj*, n = 11 CEP *Δpige*). B) B6 mice given a primary infection with CEP *hxgprt-* and 35 days later (i.e. “chronically infected”) were given a secondary infection with 5×10^4^ parasites of the indicated variants of the type I RH strain and tracked for survival. Cumulative survival from 4 experiments is plotted (mice; n = 9 RH, n = 12 RH *Δpigj*, n = 6 RH *Δpige*). C) A/J mice were given primary infections of 10^4^ parasites of either CEP or CEP *Δpigj* and monitored for survival for 30 days. Cumulative survival from 2 experiments is plotted (mice; n = 20 CEP, n = 20 CEP *Δpigj*,) D) After A/J mice were given a primary infection with the avirulent type III strain CEP, 35 days later were given a secondary infection with 5×10^4^ parasites of the indicated type I strains and tracked for survival. A single experiment was performed (mice; n = 5 RH, n = 3 RH *Δpigj*, n = 2 GT1, n = 5 GT1 *Δpigj*, n = 10 GT1 *Δpige*). For survival analysis, significance was determined by Log-rank (Mantel-Cox) test, **** p <0.001, *** p<0.01.

### Complementation of *PIGJ* partially rescues the survival phenotype

Given the most consistent phenotype surrounding the GPI sidechain occurred with the CEP *Δpigj* strain, a PIGJ complementation strain was generated in this background and efforts were focused on primary infection virulence. The full-length coding region with introns spliced out of *TgVEG_207750* was amplified from cDNA and fused with 1000 bps of the gene’s promoter, followed by delivery as a transgene into CEP *Δpigj* parasites using a pLIC-DHFR-3xHA plasmid which adds a C-terminal hemagglutinin tag to the gene of interest (54). Western blot analysis shows the C2 complementation clone, CEP *Δpigj* + *PIGJ_3xHA_*, has its GPI sidechain glycoforms restored albeit to levels lower than wildtype (**Fig 5A**), and the presence of the HA tag is detectable by fluorescence microscopy (**Fig 5B**). Both diffuse and puncta distribution of PIGJ_3xHA_ are seen throughout the cell of the parasite with no distinctive localization pattern to determine its subcellular localization. Survival, weight loss and cyst formation following primary infection with the C2 complementation strain were assessed. While complete rescue of the avirulence phenotype was not attained following C2 infections, a significant increase in mouse survival and reduced weight loss occurred during primary infections (**Fig 5C**), indicating a partial complementation of the virulence phenotype is achieved. When brain cyst burdens were analyzed in the survivors of the primary infections, significantly more brain cysts were detected in the mice infected with CEP *Δpigj* compared to wildtype strains (**Fig 5D**). However, the complementation strain was not significantly different compared to the WT or *Δpigj* mutant strains (**Fig 5D**).

**Figure 5:**
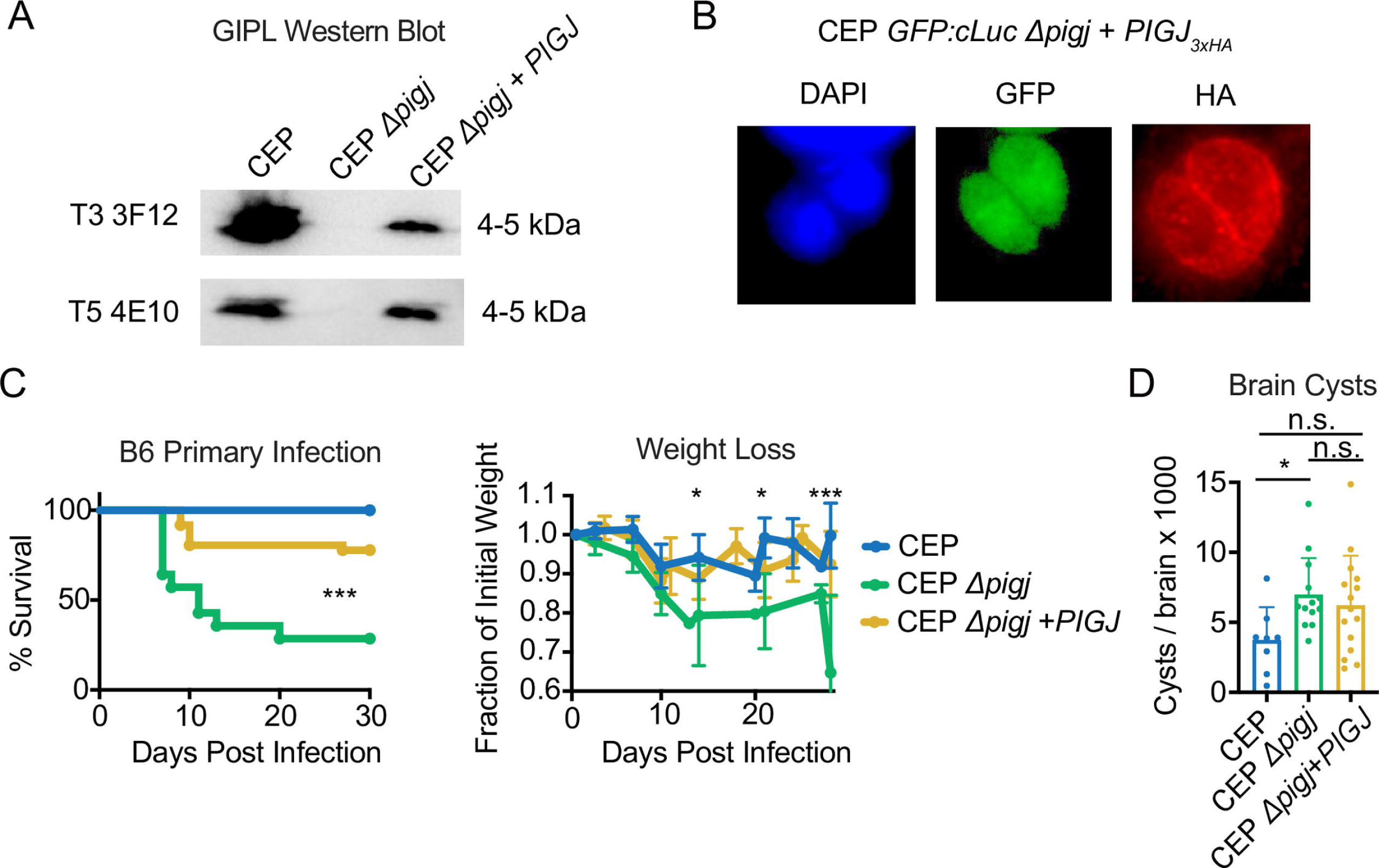
Complementation of PIGJ into *Δpigj* partially restores avirulence of the type III CEP strain. A) Western blot analysis using glycoform specific antibodies for the two GPI sidechains demonstrates GIPL reactivity is restored to the complementation strain. B) Fluorescence microscopy showing HA-tagged PIGJ localization in CEP Δ*pigj* + *PIGJ_3XHA_* parasites. The complementation strain expresses GFP, as all CEP GPI mutants were derived from a CEP *hxgprt-GFP:cLuc* parental strain. C) Mice were injected i.p. with 10^4^ parasites of CEP, CEP *Δpigj*, or CEP Δ*pigj* + *PIGJ* strains and monitored for survival for 30 days. Weight loss after primary infection was normalized to the initial weight (=1), the average +/- SD of each cohort is shown. Cumulative results from 5 experiments are plotted (mice; n = 4 CEP, n = 14 CEP *Δpigj*, n = 36 CEP *Δpigj + PIGJ*). D) Brain cysts were numerated from survivors of the indicated. Each dot represents an individual mouse, and the cumulative average +SD from 2 experiments is plotted. Statistical significance for weight loss was calculated by one-way ANOVA with multiple comparisons and a Dunnett correction (D14 and D21 * p<0.05 CEP vs. CEP *Δpigj*, D28 *** p<0.001 CEP vs. CEP *Δpigj*). Statistical significance for brain cysts was calculated by a one-way ANOVA with multiple comparisons and a Tukey correction; * p<0.05, n.s. non-significant.

To assess whether fitness defects may explain the *in vivo* phenotypes, lytic cycle analysis was performed. Consistent with genome wide fitness studies (4) (**Fig 1A**), deletion of PIGJ did not impede parasite growth in HFFs or MEFs, nor attachment to host cells (**Fig S9**). It was noted CEP *Δpigj* had significantly larger plaque areas than wildtype CEP, and this gain in plaque size was retained in the complementation strain, suggesting that either the level of complementation of PIGJ was insufficient or the background of the *Δpigj* strain gained a growth advantage during cloning irrespective of the *PIGJ* gene (**Fig S9**). Regardless, differences in *in vitro* growth rates between strains do not correlate with their pathogenesis phenotypes. In summary, virulence inversely correlates with the expression of *PIGJ* and implicates the GPI sidechain of *T. gondii* as a factor that inhibits pathogenesis.

### Early parasite burden and cytokine responses are unchanged following CEP *Δpigj* primary infections

The early weight loss and death during primary infections pointed to a possible immune defect responsible for the increased virulence of the PIGJ mutants. Therefore, mice were infected with either the wildtype CEP or CEP *Δpigj* strains and multiple organs were assessed for parasite burden as well as cytokine responses in the peritoneal cavity (**Fig 6**). *T. gondii* elicits a robust TH1 response and IFNγ is essential for survival, therefore gene expression of major cytokines known to be involved in TH1 immunity (*Il6, Il12b, Ifng, Cxcl10)* and its regulation (*Il10*) were measured. As GIPL induces robust TLR2/4-dependent TNFα production in macrophages (16), this cytokine was measured (*Tnfa*), as well as type I IFN (*Ifna, Ifnb1*) and type I IFN induced genes (*Isg15, Mx1*) due to the protective role this pathway plays during primary *T. gondii* infection (55). While there were slight increases in some cytokine response to the PIGJ mutant compared to wildtype infections, no significant differences were detected (**Fig 6A, C, E**). Moreover, parasite burden in the peritoneum, spleen, lung, and liver appeared the same between mutant and parental strain as measured by qPCR (**Fig 6B, D, F**), or by flowcytometry for GFP+ parasites in the peritoneum (**Fig S10**). Therefore, host susceptibility to primary infection with the PIGJ mutant is not related to overt dysregulation in the cytokine response or parasite burden during acute infection.

**Figure 6:**
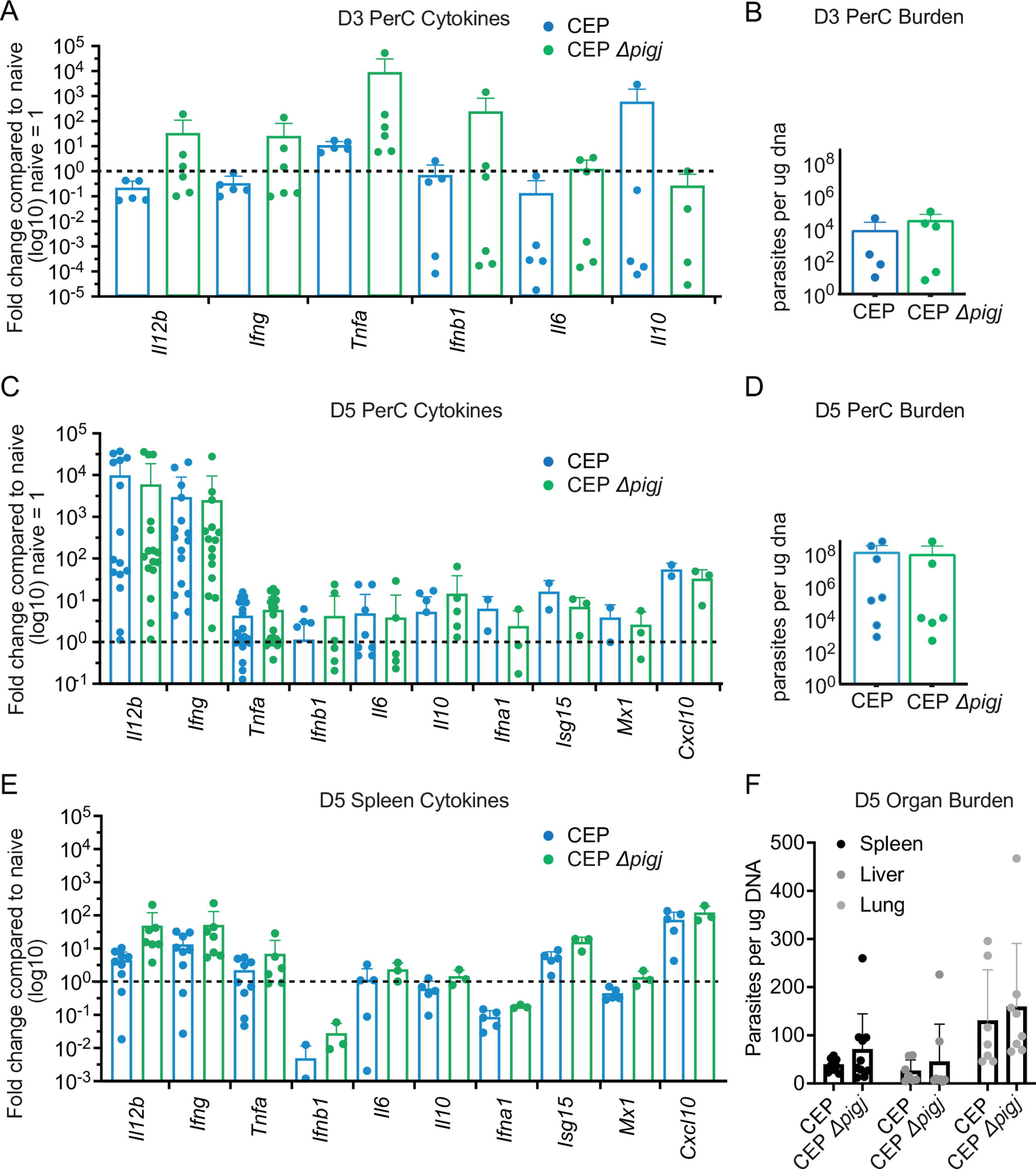
Early parasite burden and cytokine responses are similar between PIGJ mutant and wildtype strains. A) Day 3 and C) day 5 after infection with the indicated parasite strains, peritoneal cavity exudate cells and E) splenocytes were harvested and RNA isolated. cDNA was synthesized from RNA and qPCR was performed to measure gene expression levels, which were normalized to uninfected mouse levels (naive=1). Cumulative results from 2-5 experiments are plotted. Each dot is the result of an individual mouse. B) Day 3 and D) 5 after infection, DNA was isolated from the peritoneal exudate cells and qPCR was performed using the *Toxoplasma* specific B1 primers to measure parasite burden relative to the DNA in each sample. Cumulative results of 2-3 experiments is plotted, with each dot representing an individual mouse. F) 5 days after infection, spleen, liver, and lungs were harvested, DNA was isolated, and qPCR was performed using the *Toxoplasma* specific B1 primers to measure parasite burden between infections. Each dot represents the result of an individual mouse and cumulative results from 2 experiments are plotted. Statistics were performed with an unpaired t-tests, for which no condition revealed a significant difference between CEP and CEP *Δpigj* infections.

### Antibody reactivity, function and expression of GPI-anchored SAGs remain largely intact to PIGJ mutants

We wondered whether the loss of the sidechain influenced regulation of GPI anchored protein expression and/or antibody recognition of GPI-APs, and if this correlated with parasite virulence. Some reports suggest GPI-sidechains may alter the protein conformation of the GPI-AP (56), and the amount of time a GPI-AP spends folded over on the plasma membrane (57), both of which could potentially impact antibody recognition of GPI-APs. To this end, the surface expression of major GPI anchored surface proteins SAG1, SAG3, and p35 on wildtype and *Δpigj* strains was measured with monoclonal antibodies by flow cytometry. For both RH and CEP strains, deletion of PIGJ did not prevent surface expression nor detection of the aforementioned SAGs (**Fig 7A**).

**Figure 7:**
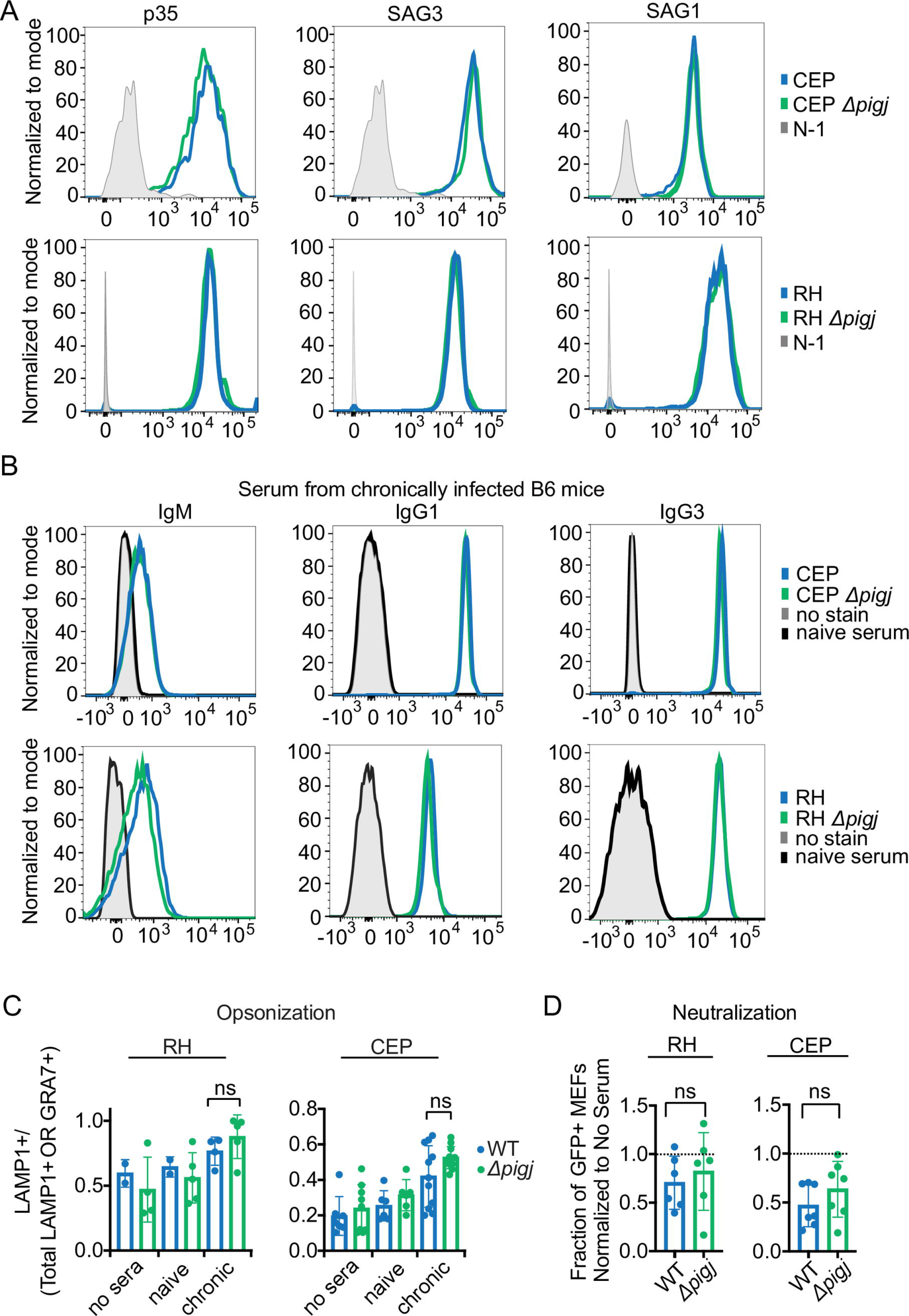
Surface expression of GPI-anchored SAGs, antibody recognition and functions against PIGJ mutants are intact. A) Parasite surface expression of p35, SAG3, and SAG1. Fixed parasites were incubated with primary antibodies against the respective surface antigen, and secondary fluorescent anti-isotype antibodies were used to measure via flow cytometry. Representative histograms of 4 different experiments shown. CEP and CEP *Δpigj* (top row); RH and RH *Δpigj* (bottom row). B) Fixed parasites were incubated with serum from CEP chronically infected C57BL/6J mice, and antibodies bound to parasites were detected with fluorescent anti-isotype antibodies. Representative histograms of 2-3 experiments displaying parasite-specific antibody reactivity to the indicated CEP strains at 10^-2^ serum dilution (top row), or RH strains (bottom row). C) Parasites were incubated with 1% serum from CEP chronically infected C57BL/6J mice (“chronic serum”) or naïve mice for 20 minutes before allowing to invade or be phagocytosed for 40 minutes. Opsonization was calculated as LAMP1+/ total LAMP1+ or GRA7+ for each parasite observed. Each dot represents the ratio obtained after counting 100 parasites by fluorescence microscopy for an individual serum, and samples were blinded. Plotted is the average ratio +SD; no serum controls were also assessed. D) Parasites were incubated with 10% chronic serum from C57BL/6J mice for 20 minutes before allowing to invade MEFs for 2 hours. MEFs were measured for GFP+ (CEP strains) or by intracellular staining with FITC-labeled parasite-specific antibodies (RH strains) indicating parasite invasion and normalized to infections in the absence of serum. Each dot represents the result from an individual serum. Statistics for opsonization was calculated with one-way ANOVA with Tukey correction and neutralization were calculated with unpaired t-tests; ns, non-significant

Parasite-specific antibody derived from the sera of mice infected with *T. gondii* allows a global assessment for any defects that polyclonal antibody recognition might have against a multitude of surface antigens regulated by the GPI sidechain. To this end, parasites were incubated with serum from chronically infected mice, and fluorescently labeled anti-isotype secondary antibodies were used to detect isotypes of parasite-bound serum antibodies as previously described (53). Again, no differences were observed for IgM, IgG3, IgG1, Ig2a/c and IgG2b reactivity to wildtype and PIGJ mutants over a range of serum dilutions (**Fig 7B**) (not shown). Complement component 3 (C3) binds the surface of *T. gondii* and is required for host resistance to infection (58). Incubating parasites with naïve serum as a source of complement and detection with an anti-C3 antibody revealed no difference in C3 recognition between WT and PIGJ mutant strains (**Fig S11**).

To screen for any functional defects of immune or naïve sera due to the loss of the GPI sidechain, opsonization and neuralization assays were performed. For opsonization, parasites were incubated with sera before being added to bone marrow derived macrophages (BMDMs) and allowed to either invade or be phagocytosed. Via microscopy, successful invasion, and formation of the parasitophorous vacuole is marked by GRA7+, while LAMP-1 marks the phagolysosome and when associated with parasites, indicates opsonization (59). No significant differences in opsonization were detected between parental and GPI mutant strains, although the CEP *Δpigj* strain trended slightly higher compared to the parental CEP strain (**Fig 7C**). To assay neutralization, serum coated parasites were incubated with mouse embryonic fibroblasts (MEFs) and assessed for relative invasion by flow cytometry, and no differences in antibody neutralization were observed between strains (**Fig 7D**). Therefore, effector functions of immune and naïve sera are largely intact to GPI sidechain null *T. gondii* parasites.

### The terminal glucose is required for immune serum IgM recognition of GIPL

Given the terminal Glc but not the GalNAc+Glc GPI sidechains is recognized by serum from humans latently infected with *T. gondii* (21), GIPL reactivity of serum from chronically infected mice was assessed against the GPI mutants by western analysis. Sera from both C57BL/6J (**Fig 8A**) and A/J mice were used (**Fig 8B**). First, GIPL reactivity, as observed by a band at 4-5 kDa when probed with anti-mouse IgM, is more robust using A/J mouse serum, but faint recognition can be seen with the C57BL/6J serum as well. Second, and consistent with the flow cytometry data, antibody reactivity to parasite antigens 18 kDa and larger did not change between the wildtype and knockout parasites. Third, RH GIPL is not detected by poly clonal immune sera, which is again consistent with previous mass-spectrometry and western blot analysis. Importantly, loss of IgM GIPL reactivity to *Δpige* and *Δpigj* was clearly observed for both GT1 and CEP mutants (**Fig 8A B**). Therefore, the GalNAc + Glc glycoform of the GPI sidechain is required for serum antibody recognition of GIPL, as previously inferred (17,21). However, given the drop in serum IgM reactivity to both virulent (*Δpigj*) and avirulent (*Δpige*) mutants, we reason this phenotype is unrelated to primary and secondary infection virulence in mice.

**Figure 8:**
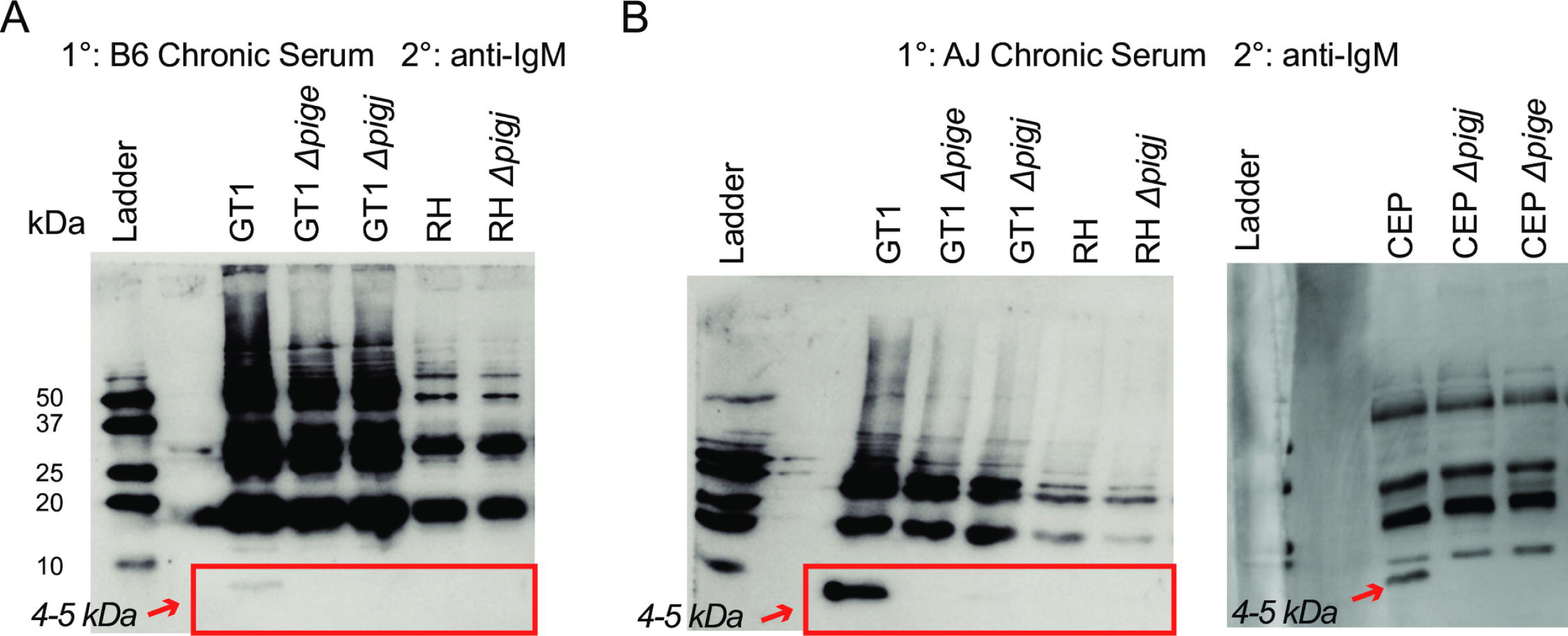
Sera IgM recognition of GIPL requires the terminal glucose. Parasite lysate was analyzed via western blot. Blots were probed with serum from CEP chronically infected A) C57BL/6J (B6) or B) A/J mice and anti-mouse IgM-HRP was used to detect IgM reactivity to parasite lysate antigens of the indicated genotypes. Red boxes and arrows indicate the region where the low molecular weight 4-5 kDa GIPL antigen migrates.

### Non-significant macrophage tropism but enhanced CD36 binding of PIGJ mutants

To test whether cellular tropism of GFP+ expressing parasites may explain difference in host susceptibility early in infection, mice were infected with 10^6^ parasites and 3 hours later peritoneal exudate cells were analyzed by flow cytometry as previously described (60). Invasion of PECs was estimated by analyzing FSC and SSC parameters of GFP+ events to differentiate by size (**Fig 9A**). Given similar GFP signal (**Fig 9B**), no differences in cell associated and intracellular (FSC^int-hi^ SSC^int-hi^) vs. non-cell associated and extracellular (FSC^lo^ SSC^lo^) GFP+ parasites were observed between strains (**Fig 9C**).

**Figure 9:**
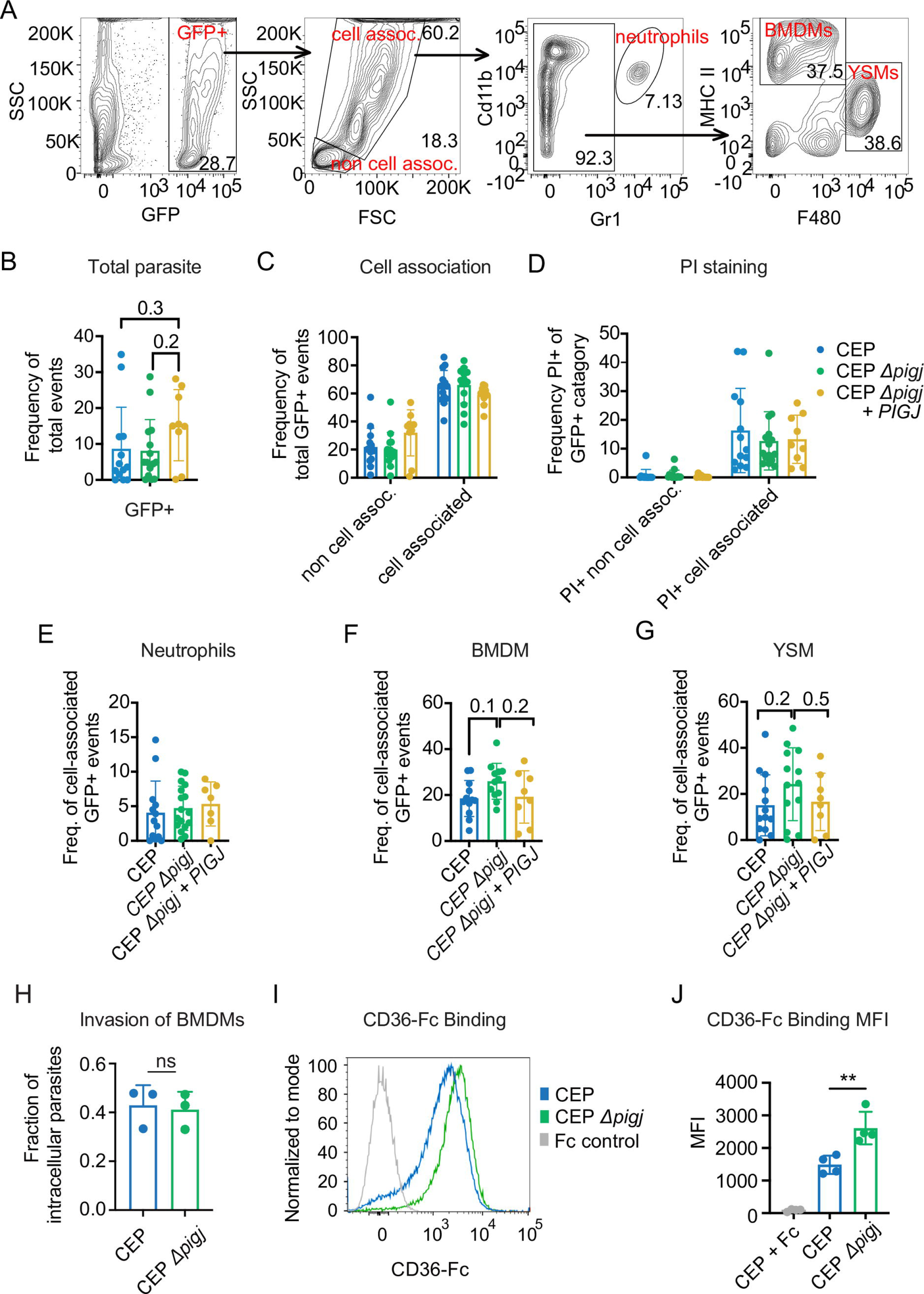
Enhanced CD36 binding and slight macrophage tropism 3 hours after infection. Mice (C57BL/6J) were injected i.p. with 10^6^ parasites of either CEP, CEP *Δpigj* or CEP *Δpigj* + *PIGJ* and PECs were analyzed 3 hours later by flow cytometry. A) Gating strategy for B-G. B) Total GFP+ signal in the peritoneal exudate; GFP frequency of total events. C) GFP+ events were separated by size using FSC and SSC parameters to determine extracellular or “non cell associated” and intracellular or “cell associated” GFP+ parasites. D) Frequency PI+ cells of the indicated GFP+ category. E) Frequency of cell associated GFP+ events that were GR-1^hi^ Cd11b+ neutrophils. F) Frequency of cell associated GFP+ events that were MHCII^hi^ F4/80^lo^ bone marrow-derived macrophages (BMDM), or G) MHCII^lo^ F4/80^hi^ yolk sack-derived macrophages (YSM). Plotted is cumulative of 7 experiments, each dot represents a single mouse (mice; n = 12 wt, n = 12 *Δpigj*, n = 8 *Δpigj +PIGJ*). Statistics performed were one-way ANOVAs with multiple comparisons and a Tukey correction; none were significant, values are indicated. H) *In vitro* differentiated BMDMs were plated on coverslips overnight and infected with 100 and 300 of GFP+ parasites per well, and parasites were allowed 20 minutes to invade cells before being fixed and stained for fluorescence microscopy. No permeabilization was used, and SAG1 staining was performed. Intracellular parasites were not stained with SAG1 but were GFP+, while extracellular parasites are SAG1+GFP+. The fraction of intracellular parasites is plotted, each dot is a representative of one experiment, n = 3 experiments. I) Parasites were incubated for 1 hr with recombinant CD36-Fc and CD36 binding was measured with anti-human-IgG-Daylight-550 via flow cytometry. Representative histogram, and J) average +SD MFI of CD36 binding from 4 experiments is plotted.

Cell death, as inferred by PI staining, was also similar between parasites, suggesting similar death kinetics of parasite infected peritoneal cells (**Fig 9D**). The cell associated GFP+ events were further analyzed for markers to identify infected cell types as a means of assessing parasite tropism. Gr-1^hi^ Cd11b+ neutrophils had no significant differences in parasite infection (**Fig 9E**). Regarding macrophages, there was a higher frequency of bone marrow-derived macrophages (MHCII^hi^ F4/80^lo^ “BMDM” cells) infected with CEP *Δpigj* compared to WT strains, indicating a tropism for BMDMs (**Fig 9F**). Infection frequencies of BMDMs with the complementation strain mirrored that of the WT strain and was reduced in relation to *Δpigj*. A similar trend was observed for MHCII^hi^, F4/80^hi^ yolk-sack macrophages (YSM) (**Fig 9G**). However, due to the biological variability, differences in BMDM or YSM tropism between strains did not reach statistical significance. To test whether this subtle tropism was related to enhanced invasion of macrophages, invasion assays were performed using *in vitro* differentiated BMDMs. When BMDMs were infected for 20 minutes and intracellular/extracellular parasites were quantified, no difference in cellular invasion was observed between the parental and *Δpigj* strains (**Fig 9H**).

CD36 is a scavenger receptor expressed in many tissues and immune cells that plays a role in fatty acid uptake, angiogenesis and phagocytosis (61). Moreover, *T. gondii* tropism for macrophages is mediated in part by CD36 (60). Therefore, parasites were incubated with recombinant CD36, and binding was measured via flow cytometry. CD36 binding was found to be significantly increased to CEP *Δpigj* compared to wildtype CEP strains (**Fig 9I J**). Whether macrophage tropism is inhibited by PIGJ, presumably through the addition of the GPI sidechain, and whether strain-dependent differences in macrophage association are mediated by CD36 is unknown.

### Virulence of *Δpigj* strains is galectin-3 and sex dependent, but not TLR-2 nor -4 dependent

Since TLR2 and TLR4 (16) and galectin-3 recognize *T. gondii* GIPL (50), we hypothesized that TLR and/or galectin-3 signaling would mediate differences in the PIGJ-dependent virulence. First, *Tlr2/Tlr4* -/- double knockout mice were used to study survival. However, the double knockout mice similarly succumbed to CEP *Δpigj* while surviving to CEP (**Fig 10A**), suggesting that differences in the GPI sidechain do not impact host-parasite interactions mediated by these PRRs. In contrast, mice lacking the gene for galectin-3, *Lgals3 -/-*, survived both to CEP and to CEP *Δpigj* infections (**Fig 10B**). This indicates that the *Δpigj* virulence observed in wildtype mice is dependent on galectin-3. The CEP *Δpigj* infected mice still had significant weight loss, but this occurred later in the infection (**Fig 10C**). Despite the weight loss, the mice survived, and did not have significant differences in brain cyst burdens (**Fig 10D**). When recombinant galectin-3 binding was measured on whole parasites via flow cytometry, an increase in galectin-3 binding to CEP *Δpigj* parasites was observed (**Fig 10E F**). It is unknown whether host factors like CD36 and galectin-3, unimpeded by a sidechain, preferentially bind the GPI mannose core to regulate the pathogenesis of *Δpigj* strains. Regardless, when mice lack galectin-3, *Δpigj* virulence is disrupted and the *Lgals3 -/-* mice are protected.

**Figure 10:**
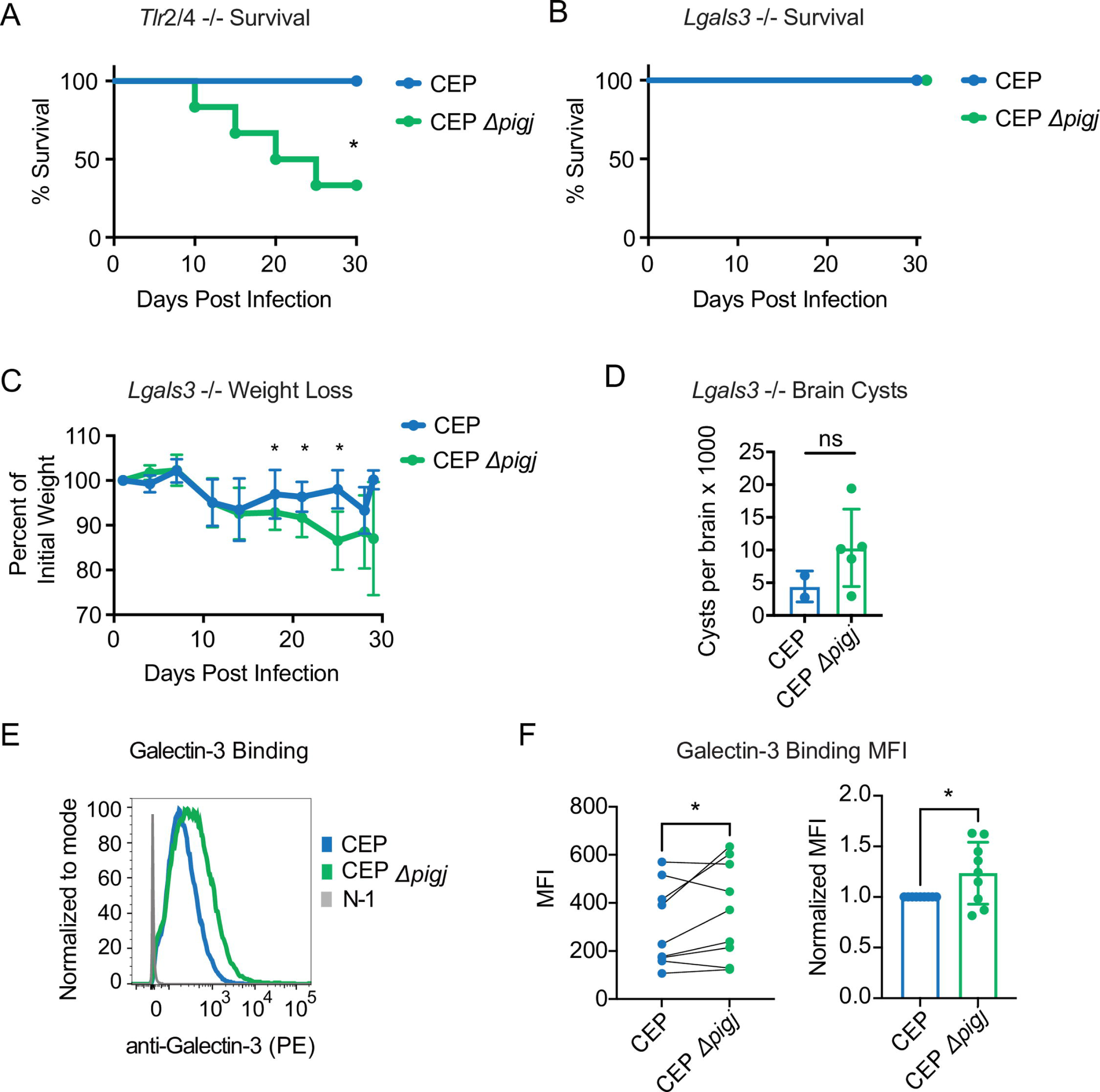
CEP *Δpigj* virulence is galectin-3 dependent. A) *Tlr2/4 -/-* mice were given primary infections of 10^4^ parasites of CEP or CEP *Δpigj* and monitored for survival for 30 days. Plotted are cumulative results from 2 experiments (mice; n = 5 CEP, n = 6 CEP *Δpigj*). B) *Lgals3 -/-* mice were given primary infections of 10^4^ parasites of CEP or CEP *Δpigj* and monitored for survival for 30 days. Plotted are the cumulative results from 4 experiments (mice; n = 15 CEP, n = 23 CEP *Δpigj*). C) Plotted is the average +/-SD fraction of initial weight for each cohort analyzed in B. D) Brain cysts were quantified from surviving mice in B. Shown is 2 experiments, (mice; n = 2 CEP, n = 5 CEP *Δpigj*). For survival analysis, significance was determined by Log-rank (Mantel-Cox) test, * p <0.05. For weight loss, significance was determined by one-way ANOVA test, * p <0.05. For brain cysts unpaired t-tests were performed, ns = non-significant. E) Whole parasites were incubated with 10 ug/mL of recombinant mouse galectin-3 and galectin-3 binding was detected with anti-Gal-3-PE by flow cytometry. Representative histogram of galectin-3 binding; N-1 control is the staining in the absence of galectin-3. F) MFIs of galectin-3 parasite binding from 9 experiments are plotted. MFI values are connected by experiment and statistical analysis was performed with a paired, one-tailed t-test. Normalized MFIs values (CEP=1) are also plotted, and statistical analysis was performed with an unpaired, one-tailed t-test; * p <; 0.05.

One final and surprising observation was that male C57BL/6J mice infected in these studies were resistant to both primary and secondary infections with *Δpigj* mutants (**Fig S12**). These mice sustained their weight similarly to the wildtype infected mice (**Fig S12B D**) and had similar survival (**Fig S12A C**). There are sex differences in a variety of immune responses, which have been reviewed (62). Specifically in *T. gondii*, females have been shown to be more susceptible to acute infections compared to males, with different cytokine kinetics, and the females were found to have higher brain cyst burdens (63). While the sex basis for this phenotype was not explored further, this observation may help unravel the mechanism behind *Δpigj* virulence.

## DISCUSSION

This study aimed to address the following questions and gaps in the field’s knowledge: 1) What are the GPI sidechain-modifying enzymes of *T. gondii*, and 2) Does the sidechain of *T. gondii* play a role in parasite virulence? In our attempts to answer these questions we made several critical observations. Firstly, we identified the two glycosyltransferases responsible for GPI sidechain modification in *T. gondii*, *Tg_207750*, named PIGJ, which adds the GalNAc to the mannose of the core backbone, and *Tg_266320*, named PIGE, which adds the glucose to the GalNAc sidechain. We confirmed their activity using mass spectrometry analysis of the GPI in the knockout parasites, which clarified ambiguities using monoclonal antibodies which apparently have some cross-reactivity to unknown epitopes. In addition to the characterization of these glycosyltransferases, we explored their role in parasite virulence in both primary and secondary infections which revealed that PIGJ mutants are virulent in both settings. This study is a first analysis for the role of the GPI sidechain for any microbe in pathogenesis.

Removal of the core GPI synthesizing enzymes is lethal in mammals and in eukaryotic microbes. However, it remained an outstanding question whether complete removal of the GPI sidechain would have fitness bearing impacts in parasites. The genes identified here do not have deleterious impacts on the lytic cycle, and the fitness scores for PIGJ and PIGE align with this supposition (**Fig 1A**). In performing reciprocal blasts using *T. gondii* PIGJ and PIGE, we find evidence that homologous enzymes exist in coccidian species, but not in more distantly related apicomplexan species such as *Plasmodium* and *Cryptosporidium* (not shown). Coccidia parasites are orally acquired parasites of warm-blooded animals that have an intra-epithelial stage in their development, but many form tissue cysts, which require parasite dissemination following infection. Whether all coccidia parasites express similar GalNAc + Glc bearing sidechains is unknown but would be expected from the identification of PIGJ and PIGE homologs across this subclass of parasites. It is tempting to speculate that the GPI sidechain, with a structure similar to that of mammals (**Fig S1**), use this similarity as the result of convergent evolution, to mimic a fundamental biological process mediated by this glycoform and required in these environments.

In our search for PIGJ and PIGE functions, we characterized and knocked out the enzymes responsible for GPI sidechain synthesis in both type I (GT1, RH) and a type III (CEP) strains of *T. gondii*. Parasites lacking PIGJ, but not those lacking PIGE, exhibit increased virulence in primary and secondary infections, demonstrating the GPI sidechain modulates parasite virulence by preventing lethality in its host. To elucidate the mechanism for increased virulence of the PIGJ mutant we characterized a variety of immune parameters and parasite burden following infection. Our findings indicate early parasite burden in the peritoneal cavity, liver, and lung are similar, yet the PIGJ mutants can evade clearance and establish higher cyst burdens in the brain. Cytokine responses, antibody reactivity, opsonization, neutralization, and parasite surface antigen expression were all similar between strains that express a GPI sidechain and those that do not. We speculate that a breakdown of infectious tolerance occurs early during infection when the parasite lacks a GPI sidechain. Whether enhanced tissue damage results following infection with GPI sidechain null parasites, is unknown.

To further explore where the pathogenesis might take place, a subtle cell tropism for BMDMs was observed in the peritoneal cavity 3 hrs after infection, and CD36 binding was found to be increased with CEP *Δpigj*. We also found that *Lgals3* -/- mice were protected against primary infections with CEP *Δpigj*, indicating a fundamental role of galectin-3 in the susceptibility of wildtype mice to CEP *Δpigj* infections. Galectin-3 is one of the few known host factors that bind *T. gondii* GIPL (50), and recombinant galectin-3 preferentially bound CEP *Δpigj* strains. Galectin-3 was previously shown to be required for macrophage TNFα production in response to *T. gondii*, and it was proposed that galectin-3 might act as a co-receptor that presents the GPI to TLR2 (50). Our data of PIGJ infections in double knockout mice of *Tlr2/4 -/-* indicated that TLR2/4 signaling was not involved in the virulence differences. It is likely there are other cellular functions mediated by galectin-3 binding to GIPL that do not involve TLR2/4, however, the exact mechanisms are unknown. Previous reports have shown that following *T. gondii* infection, galectin-3 is upregulated in peripheral tissues during acute infection of *T. gondii* and were susceptible to intraperitoneal infections with type II strains of *T. gondii* (64). Interestingly, *Lgals3* -/- mice had higher Th1 responses with increased IL-12p40 and IFNγ detected in the sera, as well as from cultured splenocytes after infection (64). It is possible the enhanced Th1 response underpins the resistance of *Lgals3* -/- mice to type III strains but renders them susceptible to more virulent type II strains. Regardless, further mechanistic insight is required to draw correlations between galectin-3 and the GPI sidechain, and to place this interaction as central to the disease outcome of PIGJ mutants.

Of further note on the CD36 pathway, it is possible that mice infected with *Δpigj* strains are more able to phagocytose parasites via CD36, allowing for enhanced infection and escape via the phagosome cellular-invasion route reported for less virulent parasite strains (59). *Cd36 -/-* susceptibility to *T. gondii* infections was suggested to correlate with a breakdown in tissue homeostasis following infection (60). This was revealed by monitoring serum indicators of tissue stress including GDF-15 and FGF21. GDF-15 induces anorexia by binding to the GFRAL receptor expressed by neurons of the hindbrain (65), while GFG21 is a hormone that controls energy homeostasis and adiposity (66) and is often induced in multiple tissues by nutrient starvation and endoplasmic reticulum stress (67). Given the PIGJ mutants appear normal with respect to parasite burden and the disease outcome is sex linked, we suspect the GPI sidechain promotes tissue homeostasis during infection by potentially controlling mechanisms of tolerance via CD36 interactions and/or other host pathways yet to be identified.

In our initial approach to confirm the loss of the Glc and GalNAc sidechain additions, we utilized glycoform specific antibodies T3 3F12 and T5 4E10. While complete removal of the sidechain in the *Δpigj* mutants was clearly observed through western blotting, the removal of the glucose addition was not. The T5 4E10 antibody, though it is described to be specific for the GalNAc + Glc glycoform, has considerable cross-reactivity to the GalNAc only glycoform, as seen in previous studies utilizing it (17,29). The T3 3F12 antibody clone is an IgG3 isotype, and the T5 4E10 clone is an IgM isotype. It is interesting to note, B-1 cells preferentially produce antibodies of IgM and IgG3 isotypes. B-1 cells are responsible for natural self-reactive IgM antibodies and are known to contribute to T-independent response, most of which are against non-protein antigens (68). We originally looked into the GPI as a non-protein antigen after we found that B-1 cells are important for immunity to secondary infections of T. gondii (53). With respect to the IgM response, we did note that IgM recognition of GIPL was highly sensitive to the terminal glucose (**Fig 8**), as previously observed using chemically synthesized GPI and serum from latently infected humans (21). Whether the antibody response to GIPL is from B-1 cells is unknown. Regardless, the lack of antibody reactivity to GIPLs of the PIGE and PIGJ mutants, cannot account for why these strains differ in primary and secondary infection virulence.

The analysis of the type I RH strain presents additional insights into the significance of the sidechain and its biosynthesis. Previous reports have indicated detection of RH GIPL and its sidechain (17). However, a recent report using mass spectrometry of RH GPI-AP revealed only the linear mannose core of GPI with no sidechain (36). A potential cause for this could be the lab adaptation of this strain. This strain was initially passaged in mice and then in tissue culture for over 60 years before ever being frozen down. Due to this, the RH strain has become lab adapted, and acquired some unique characteristics (45). For instance, RH has lost its ability to form orally infectious tissue cysts, grows much more rapidly than other strains, and has increased extracellular viability (69,70). We have discovered that the type I RH strain has lost its ability to evade immunological memory responses that are generated following vaccination or natural infection, which we demonstrated to occur with other type I strains like GT1 (43,53). It is likely that for whatever reason, RH has lost its expression of the GPI sidechain through this laboratory evolution. However, it was an initial report that the gene *Tg_207750* was almost 6 times more highly expressed in RH compared to GT1 that led us to investigate it as a GPI sidechain GT candidate (45). Overexpression may play a role in the loss of the sidechain, and a second possible explanation is an identified non-synonymous SNP that generates a L620R substitution uniquely in this strain relative to all other available sequences (**Fig 1B**) (not shown). This position is conserved as a hydrophobic core of this region of the enzyme that is involved in coordinating a PO_4_ group of the UDP-GalNAc substrate. Whether this acquired mutation in RH is inactivating to PIGJ, which in turn leads this strain to compensate by overexpressing the gene is unknown. Interestingly, full deletion of the gene increased virulence in secondary infections, suggesting an additional role for PIGJ that might be mediated by other regions of the protein. It is possible that PIGJ, like most other enzymes in the GPI processing pathway (49), belongs to a multiprotein complex that depends in part on the presence of the PIGJ protein, and that its absence disturbs some other aspect of GPI anchor assembly such as selectivity or efficiency.

In conclusion, this study identified and characterized both GPI sidechain modifying glycosyltransferases of *T. gondii*. In addition, for the first time in any microbe, the effects of the complete loss of the sidechain were studied for its role in pathogenicity. The results presented here indicate a fundamental role of the presence of the sidechain in host survival, and that when the sidechain of the GPI is lost, so is host resistance to *T. gondii.* Although most of the major surface antigens (SAGs) of *T. gondii* are GPI-anchored proteins and the GPI is known to be targeted by antibody responses in a variety of protozoan infections (19,26), our findings here would suggest that for *T. gondii* antigens, the sidechain presence does not impact antibody reactivity to GPI-anchored proteins. However, whatever other humoral functions are at play regarding host recognition of GIPL, they are wildly disrupted when the sidechain is lost. We suggest both a fundamental, and yet diverse role for the GPI sidechain in microbial pathogenesis. Based on our data, we propose a model in which enhanced CD36 and galectin-3 binding of PIGJ mutants lead to pathology, perhaps through differences in cellular tropism and unknown inflammatory mediators yet to be defined (**Fig S13**).

## ABBREVIATIONS

GPI: Glycosylphosphatidylinositol
MFI: mean fluorescence intensity

## ACKNOWLEDGEMENTS

Jean Françios-Dubremetz (U Montpellier) for the T5 4E10 ascites. Boris Striepen (U Penn) for technical support and guidance. John Boothroyd (Stanford) for the rabbit GRA7 polyclonal antibody. Gregory Barton (UC Berkeley) for the *Tlr2/4* -/- mice. Jeroen Saeij (UC Davis) for plasmids. Dave Gravano for assistance with flow cytometry (UC Merced, SCIF). The following monoclonal antibodies (produced in vitro) were obtained through BEI Resources, NIAID, NIH: Anti-*Toxoplasma gondii* Glycosylphosphatidylinositol Anchor, Clone T3 3F12, NR-50253; anti-*T. gondii* Surface Antigen 1, Clone T4 1E5, NR-50255, anti-*T. gondii* Surface Antigen 3, Clone T4 1F12, NR-50257, anti-*T. gondii* Surface Antigen P35, Clone T4 3F12, NR-50259. This work was supported by NIH grants 1R01 AI137126 and 1R21 AI176186 to KDCJ; 1R21 AI123161 to CW; R21 AI171670 to GY. JP was supported by the supplement 3R01 AI137126-04S1, and GC was supported by a UC LEADS fellowship.

## SUPPLEMENTAL FIGURE LEGENDS

**Figure S1. GPI sidechains differ between eukaryotic species.**

Schematic of the diversity of GPI sidechains between species (*T. gondii*, *P. falciparum*, *T. brucei* procyclic stage, *H. sapiens*). Note the conserved mannose backbone with species variability of sidechain modifications. Not all GPI glycoforms for each species are represented.

**Figure S2: Approach and PCR confirmation of *Tg_207750* disruption.**

Schematic of the CRISPR Cas9 targeted disruption of the *Tg_207750* (*PIGJ*) locus and insertion of the selectable marker *HXGPRT* in A, or *DHFR-TS* in B. C-E) PCR confirmation of disruption of the targeted locus and insertion of the selectable marker within the cut site using the primers indicated in panels A or B.

**Figure S3: Approach and PCR confirmation of *Tg_266320* disruption**.

Schematic of the CRISPR Cas9 targeted disruption of the T*GGT1_266320* (*PIGE*) locus and insertion of the selectable marker *HXGPRT* in A, or *DHFR-TS* in B. C-D) PCR confirmation of disruption of the target locus and insertion of the selectable marker within the cut site using primers in A or B. In the case of CEP *Δ266320*, the *HXGPRT* selectible marker inserted into the Cas9 cut site in exon one in reverse orientation, but not the second site in exon 3. In the case of GT1 *Δ266320*, the *DHFR-TS* selectible marker inseted in reverse orietnation in the Cas9 cut site in exon 1, while the cut site in exon 3 was repaired with a concatemerized DHFR-TS as prposed based upon the diagnostic PCR.

**Figure S4: Decreased T3 3F12 and T5 4E10 recognition of GT1 *Δpigj* mutants by flow cytometry**.

Fixed parasites of the indicated type I GT1 and RH parasite strains were stained with GPIL glycoform specific antibodies T3 3F12 (mouse IgG3) and T5 4E10 (mouse IgM). Secondary fluorescent anti-isotype antibodies were used to detect T3 3F12 and T5 4E10 binding. SAG3 positive parasites were analyzed to distinguish parasites from debris. Data is representative of 3-8 experiments. B) Representative histograms from the flow cytometry analysis comparing the parental with the mutant strains of the data shown in A. In addition, a no staining control is plotted for comparison (grey histogram).

**Figure S5. Fragmentation analysis of GPI-anchor glycans.**

Isolated primary GPI ions were selected in MS(1) as described in Figure 3 and subjected to collision-based MS(2) fragmentation during the nLC-MS run. Residual parent ions are labeled at the high m/z end, and decomposition products at lower m/z values. Green arrows trace sequential fragmentation pathways. A) H4N2 glycan from strain GT1. B) H3N2 glycan from GT1 *Δpige*. C) H3N1 from GT1-*Δpigj*.

**Figure S6: MALDI-TOF-MS analysis of GIPL preparations is consistent with PIGE and PIGJ as being the GPI sidechain glycosyl transferases in *T. gondii*.**

Glycans isolated from GPIL samples from tachyzoite stage parasites were N-acetylated and permethylated, and mixed with DHB matrix and analyzed as in Figure 2. A) Parental GT1 strain. The inset shows a sample prepared in CHCA rather than DHB matrix. See text for the basis of assignment of ions that differ by a m/z defect of -36. B) *Δpige*. C) *Δpigj*. D) Host cells (HFFs).

**Figure S7: nLC-MS analysis of GIPL preparations is consistent with PIGE and PIGJ as being the GPI sidechain glycosyl transferases in *T. gondii*.**

Isolated GIPL fractions described in Figure S6 were reanalyzed by nLC separation on a C18 column and hyphenated analysis in an Orbitrap mass spectrometer in positive ion mode. The left-hand column of panels shows base peak chromatograms (all ions m/z 500-2000) and extracted ion chromatograms (EIC) for each of the indicated targets (m/z ranges given in panels A and D). The ratio of ion intensities over the EIC m/z ranges shown for H3N1Ino, H3N2Ino, and H4N2Ino are shown relative to the most abundant ion. The most abundant ions are labeled in the base peak chromatogram. The right-hand column of panels shows representative mass spectra of the one or two most abundant ions eluting with the target and used for quantitating relative levels. Levels of hexosamers were not quantitated. Observed m/z values are in dark red, expected values are in black, and the charge state is as indicated. A) Parental GT1 strain. B) *Δpige*. C) *Δpigj*. D) Host cells (HFFs).

**Figure S8: C57BL/6J mice succumb to secondary infections with the type I GT1 GPI sidechain mutants similarly to the parental GT1 strain.**

C57BL/6J (B6) mice given a primary infection with the avirulent type III strain CEP, 35 days later were given a secondary infection with 5×10^5^ parasites of the type I strain GT1, GT1 *Δpigj* or GT1 *Δpige* strains and tracked for survival. Plotted is the result from one experiment (mice; n = 3 wt GT1, n=5 GT1 *Δpigj*, n=1 GT1 *Δpige*).

**Figure S9: PIGJ mutants have no apparent fitness defects *in vitro*.**

A) HFF cells in 24 well plates were infected with either CEP, CEP *Δpigj*, or CEP *Δpigj* + *PIGJ* and allowed to grow for 5 days. On day 5 plaque sizes were measured and plotted. Cumulative data from 4 experiments is shown, each dot represents a single plaque area.

B) MEFs were infected for 16 hours before being fixed and stained for GRA7 to mark the PV. Parasites per vacuole were quantified, counting 100 vacuoles per experiment, and fractions are plotted for each experiment. Cumulative data from 4 experiments is plotted, each dot is fraction obtained from an individual experiment. C) HFFs were infected with 200 parasites per well and allowed various timepoints to attach before being washed extensively and cultured for 5 days before quantifying plaque numbers, which were normalized to the plaque counts from wells that were not washed (=1). Cumulative average +/-SD from 4 experiments is plotted. For panels A-B, statistics performed were one-way ANOVAs with multiple comparisons, and a Holm-Sidak’s correction; **** p < 0.0001 (GFP vs KO), *** p < 0.001 (GFP vs C2); ns, not significant. For panel C, t-tests yielded non-significant values.

**Figure S10: Parasite burden is similar between wildtype and PIGJ mutant strains in peritoneal exudate cells as assessed by flow cytometry.**

3 and 5 days after infection with the indicated GFP expressing parasite strains, peritoneal cavity exudate cells (PerC) were harvested and PI negative cells were analyzed by flow cytometry. Average frequency (+SD) of infected GFP+ PerC among total PI-cells is shown for each infection. Cumulative results from 2-3 experiments is plotted, with each dot representing an individual mouse. Statistics were performed with an unpaired t-test and values indicated, no condition revealed a significant difference between CEP and CEP *Δpigj* infections.

**Figure S11: Complement C3b binding is unimpaired to PIGJ mutants.**

Live parasites were incubated with serum from naïve mice and C3b binding was measured with anti-C3b-APC. A) Representative histogram from 4 experiments each with different naïve serums is shown. Staining controls include use of heat inactivated serum and N-1, which is the staining in the absence of serum. C) Average +SD MFI of C3b binding, each dot is the value of a different serum.

**Figure S12: Male C57BL/6J mice are protected against *Δpigj* primary and secondary infections.**

A-B) Male C57BL/6J (B6) mice were given primary infections i.p. with 10^4^ parasites of either CEP or CEP *Δpigj* and monitored for survival in A, and weight loss in B, for 30 days. Cumulative results from 2 experiments are plotted (mice; n = 6 wt, n = 8 *Δpigj*). C-D) Male B6 mice were first given a primary infection of 10^4^ CEP parasites, and after 35 days were given a challenge infection of either RH or RH *Δpigj* and monitored for survival in C, and weight loss in D. Cumulative results from 2 experiments are plotted (mice; n = 2 wt, n = 4 *Δpigj*). For survival analyses, significance was determined by Log-rank (Mantel-Cox) test, and for weight loss, significance was determined by a one-way ANOVA test with multiple comparisons with Dunnett correction, none of which were found significant.

**Figure S13: Working model of *Δpigj* pathogenicity.**

With the loss of sidechain in *Δpigj* mutants we have found an increase in CD36 binding, an increase in BMDM tropism, and an increase in galectin-3 binding. We hypothesize that there are other inflammatory signals possibly leading to tissue damage that ultimately are responsible for the death of the hosts, and the cause for increased pathogenicity in sidechain-null mutants.

## SUPPLEMENTAL TABLE LEGEND

**Table S1. Parasite strains, oligos and plasmids.**

Parasite strains, oligos and plasmids used in this study are described and their origins indicated.

## Notes

### Competing Interest Statement

The authors have declared no competing interest.

### Summary of Updates

To post Supplemental Figures and Supplemental Table

