## Supplemental Figures 1-13 for "The GPI sidechain of *Toxoplasma gondii* prevents parasite pathogenesis"

### Fig S1

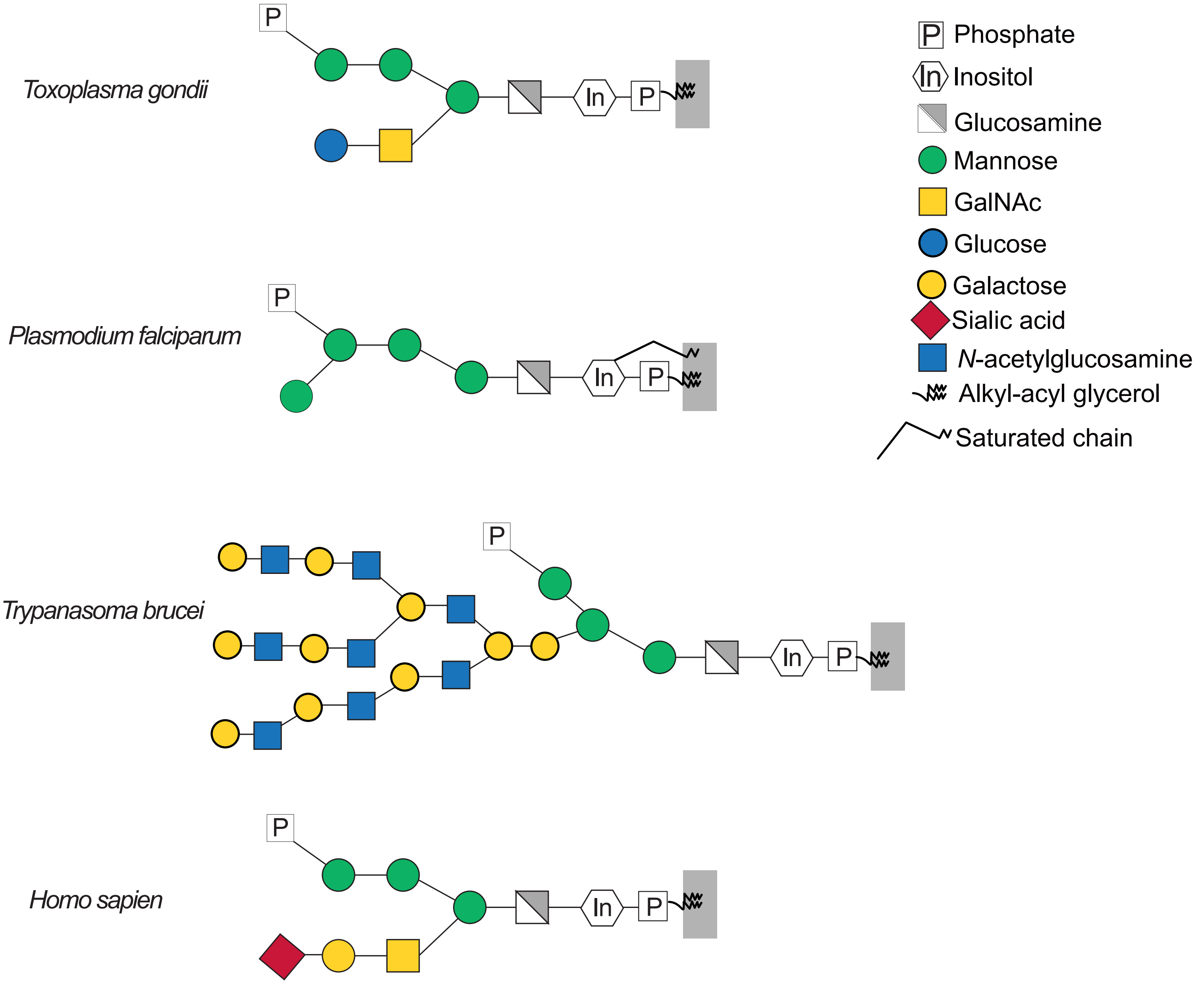
Toxoplasma gondii
Plasmodium falciparum
Trypanasoma brucei
Homo sapien
P
In
Phosphate
Inositol
Glucosamine
Mannose
GalNAc
Glucose
Galactose
Sialic acid
N-acetylglucosamine
Alkyl-acyl glycerol
Saturated chain

Fig S2

A CRISPR Cas9-targeted disruption of *Tg_207750*

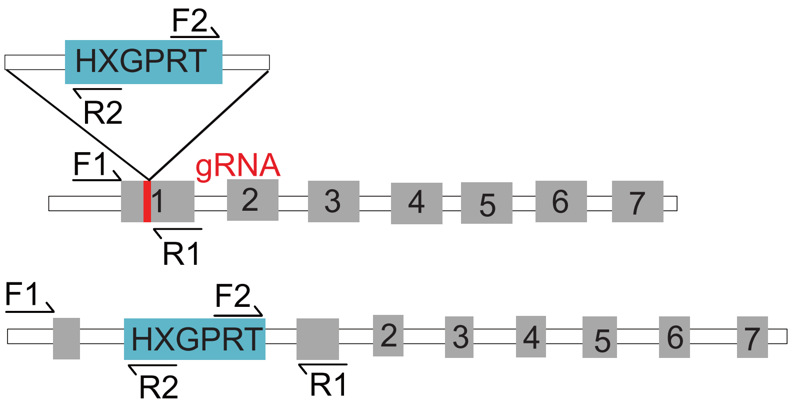


B CRISPR Cas9-targeted disruption of *TGGT1_207750*

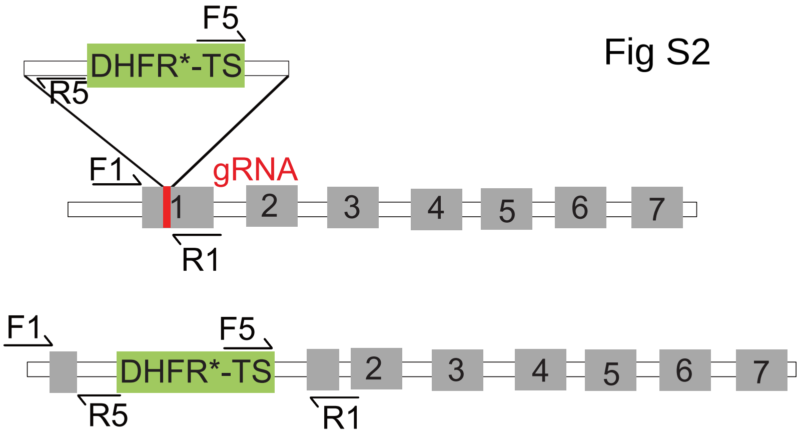


C

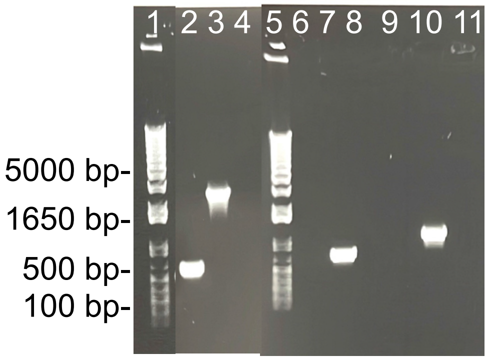


| Well | Sample | Primers |
|---|---|---|
| 1 | Ladder | |
| 2 | RH *Δku80* | F1 & R1 |
| 3 | RH *Δku80Δ207750::HXGPRT* | F1 & R1 |
| 4 | H2O | F1 & R1 |
| 5 | Ladder | |
| 6 | RH *Δku80* | F1 & R2 |
| 7 | RH *Δku80Δ207750::HXGPRT* | F1 & R2 |
| 8 | H2O | F1 & R2 |
| 9 | RH *Δku80* | F2 & R1 |
| 10 | RH *Δku80Δ207750::HXGPRT* | F2 & R1 |
| 11 | H2O | F2 & R1 |

D

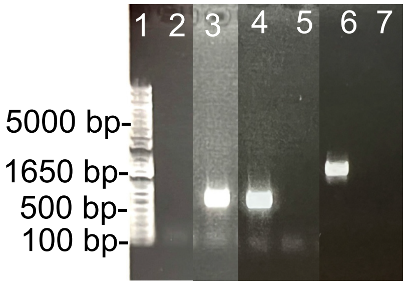


| Well | Sample | Primers |
|---|---|---|
| 1 | Ladder | |
| 2 | CEP *hpt- Δ207750* | F1 & R1 |
| 3 | CEP *hpt-* | F1 & R1 |
| 4 | CEP *hpt- Δ207750* | F1 & R2 |
| 5 | CEP *hpt-* | F1 & R2 |
| 6 | CEP *hpt- Δ207750* | F2 & R1 |
| 7 | CEP *hpt-* | F2 & R1 |

E

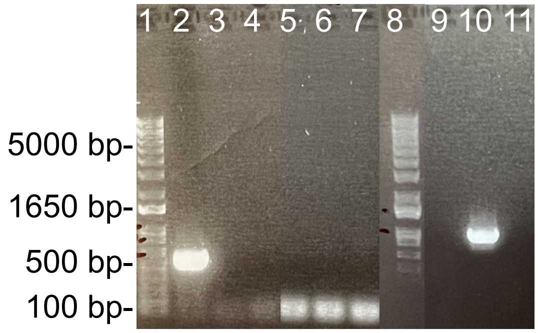


| Well | Sample | Primers |
|---|---|---|
| 1 | Ladder | |
| 2 | GT1 | F1 & R1 |
| 3 | GT1 *Δ207750::DHFR-TS* | F1 & R1 |
| 4 | H2O | F1 & R1 |
| 5 | GT1 | F1 & R5 |
| 6 | GT1 *Δ207750::DHFR-TS* | F1 & R5 |
| 7 | H2O | F1 & R5 |
| 8 | Ladder | |
| 9 | GT1 | F5 & R1 |
| 10 | GT1 *Δ207750::DHFR-TS* | F5 & R1 |
| 11 | H2O | F5 & R1 |

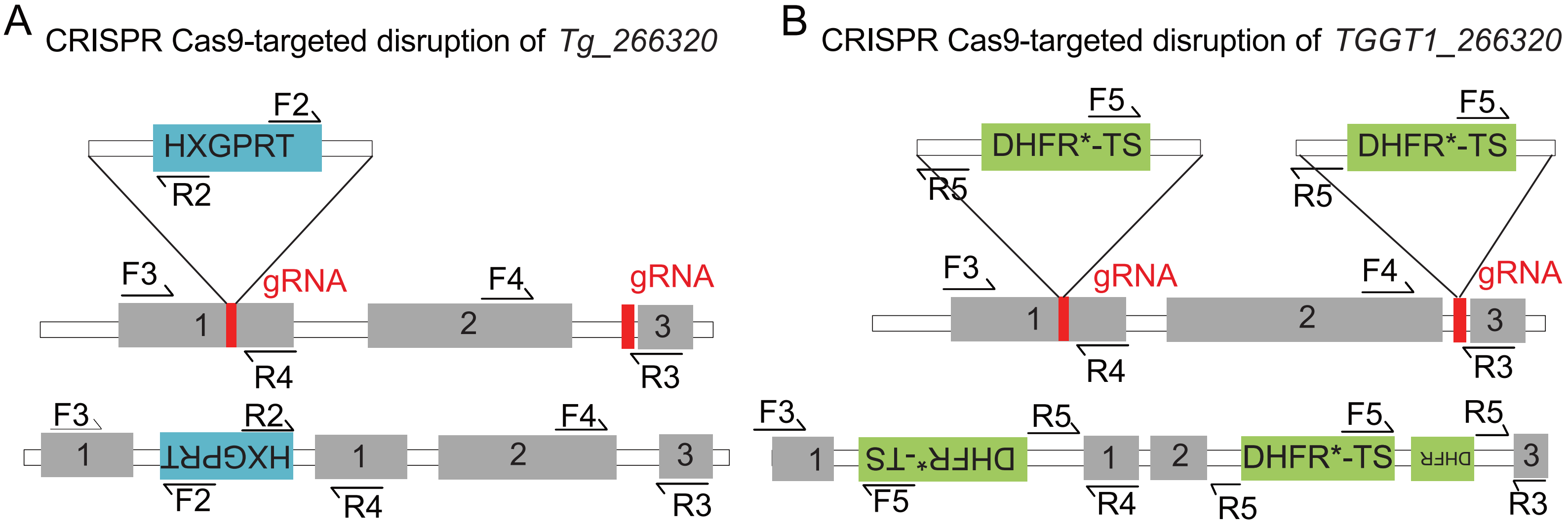


C

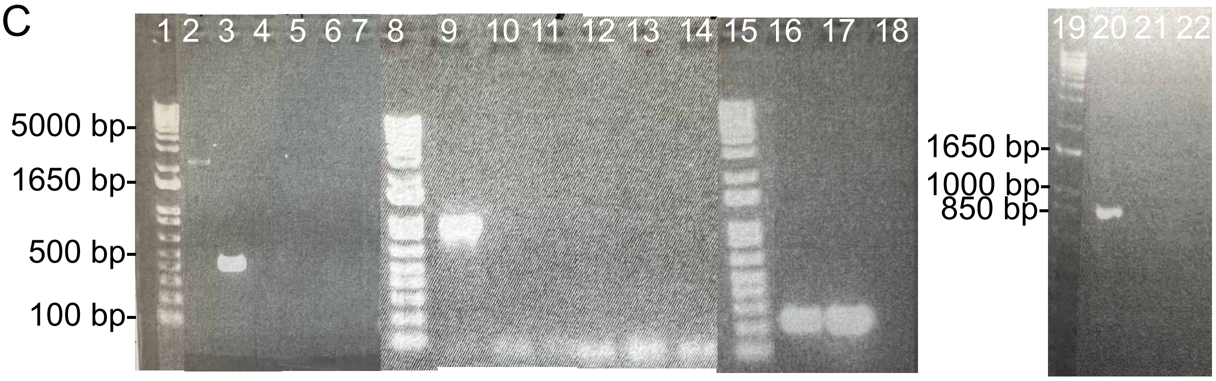


| Well | Sample | Primers | Well | Sample | Primers | Well | Sample | Primers |
|---|---|---|---|---|---|---|---|---|
| 1 | Ladder | | 8 | Ladder | | 15 | Ladder | |
| 2 | CEP *hpt- Δ266320* | F3 & R4 | 9 | CEP *hpt- Δ266320* | R4 & R2 | 16 | CEP *hpt-* | F4 & R3 |
| 3 | CEP *hpt-* | F3 & R4 | 10 | CEP *hpt-* | R4 & R2 | 17 | CEP *hpt- Δ266320* | F4 & R3 |
| 4 | H2O | F3 & R4 | 11 | H20 | R4 & R2 | 18 | H2O | F4 & R3 |
| 5 | CEP *hpt- Δ266320* | F3 & R2 | 12 | CEP *hpt-* | R4 & F2 | 19 | Ladder | |
| 6 | CEP *hpt-* | F3 & R2 | 13 | CEP *hpt- Δ266320* | R4 & F2 | 20 | CEP *hpt- Δ266320* | F3 & F2 |
| 7 | H2O | F3 & R2 | 14 | H20 | R4 & F2 | 21 | CEP *hpt-* | F3 & F2 |
| | | | | | | 22 | H2O | F3 & F2 |

D

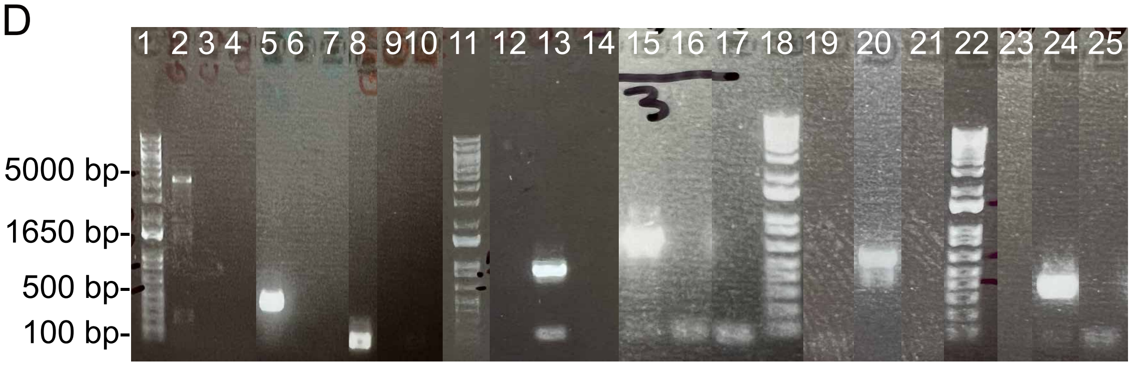


| Well | Sample | Primers | Well | Sample | Primers |
|---|---|---|---|---|---|
| 1 | Ladder | | 15 | GT1 *Δ266320::DHFR-TS* | R5 & R4 |
| 2 | GT1 | F3 & R3 | 16 | GT1 | R5 & R4 |
| 3 | GT1 *Δ266320::DHFR-TS* | F3 & R3 | 17 | H2O | R5 & R4 |
| 4 | H2O | F3 & R3 | 18 | Ladder | |
| 5 | GT1 | F3 & R4 | 19 | GT1 | F3 & F5 |
| 6 | GT1 *Δ266320::DHFR-TS* | F3 & R4 | 20 | GT1 *Δ266320::DHFR-TS* | F3 & F5 |
| 7 | H2O | F3 & R4 | 21 | H2O | F3 & F5 |
| 8 | GT1 | F4 & R3 | 22 | Ladder | |
| 9 | GT1 *Δ266320::DHFR-TS* | F4 & R3 | 23 | GT1 | R5 & R3 |
| 10 | H2O | F4 & R3 | 24 | GT1 *Δ266320::DHFR-TS* | R5 & R3 |
| 11 | Ladder | | 25 | H2O | R5 & R3 |
| 12 | GT1 | F5 & R3 | | | |
| 13 | GT1 *Δ266320::DHFR-TS* | F5 & R3 | | | |
| 14 | H2O | F5 & R3 | | | |

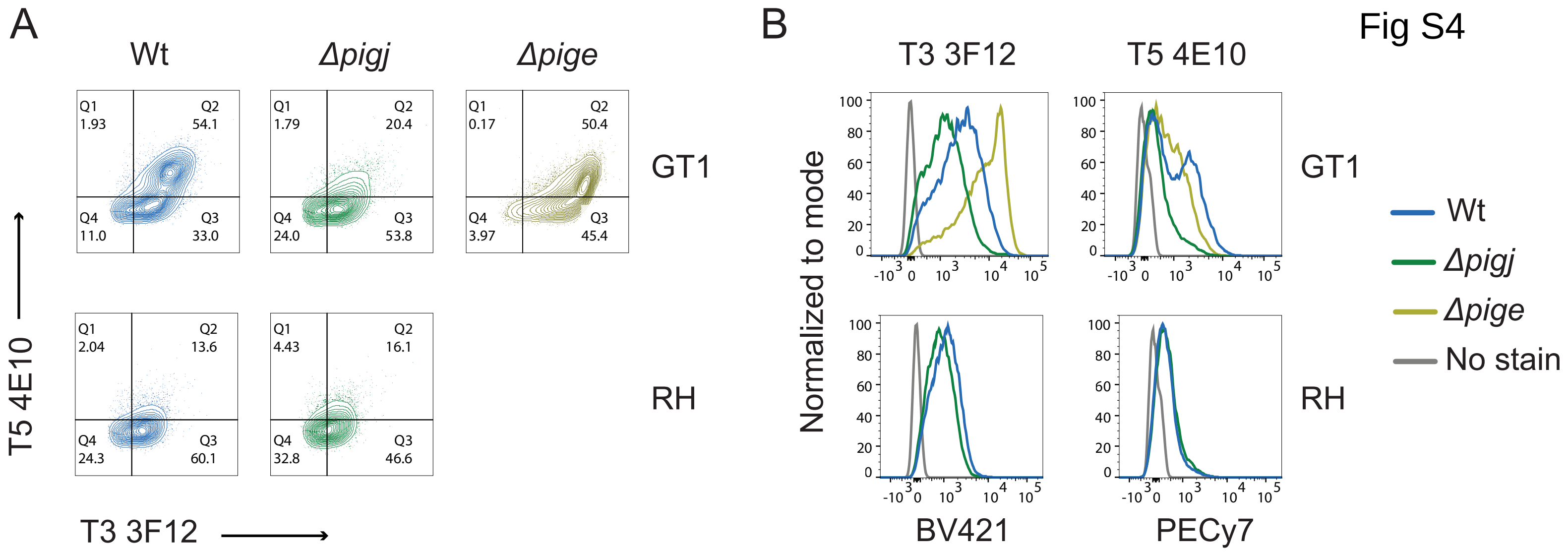
A
Wt
Δpigj
Δpige
GT1
RH
T5 4E10
T3 3F12
Q1 1.93
Q2 54.1
Q4 11.0
Q3 33.0
Q1 1.79
Q2 20.4
Q4 24.0
Q3 53.8
Q1 0.17
Q2 50.4
Q4 3.97
Q3 45.4
Q1 2.04
Q2 13.6
Q4 24.3
Q3 60.1
Q1 4.43
Q2 16.1
Q4 32.8
Q3 46.6
B
T3 3F12
T5 4E10
Normalized to mode
BV421
PECy7
Wt
Δpigj
Δpige
No stain

Fig S4

**MS(2) analysis of GPI anchors**

Fig S5

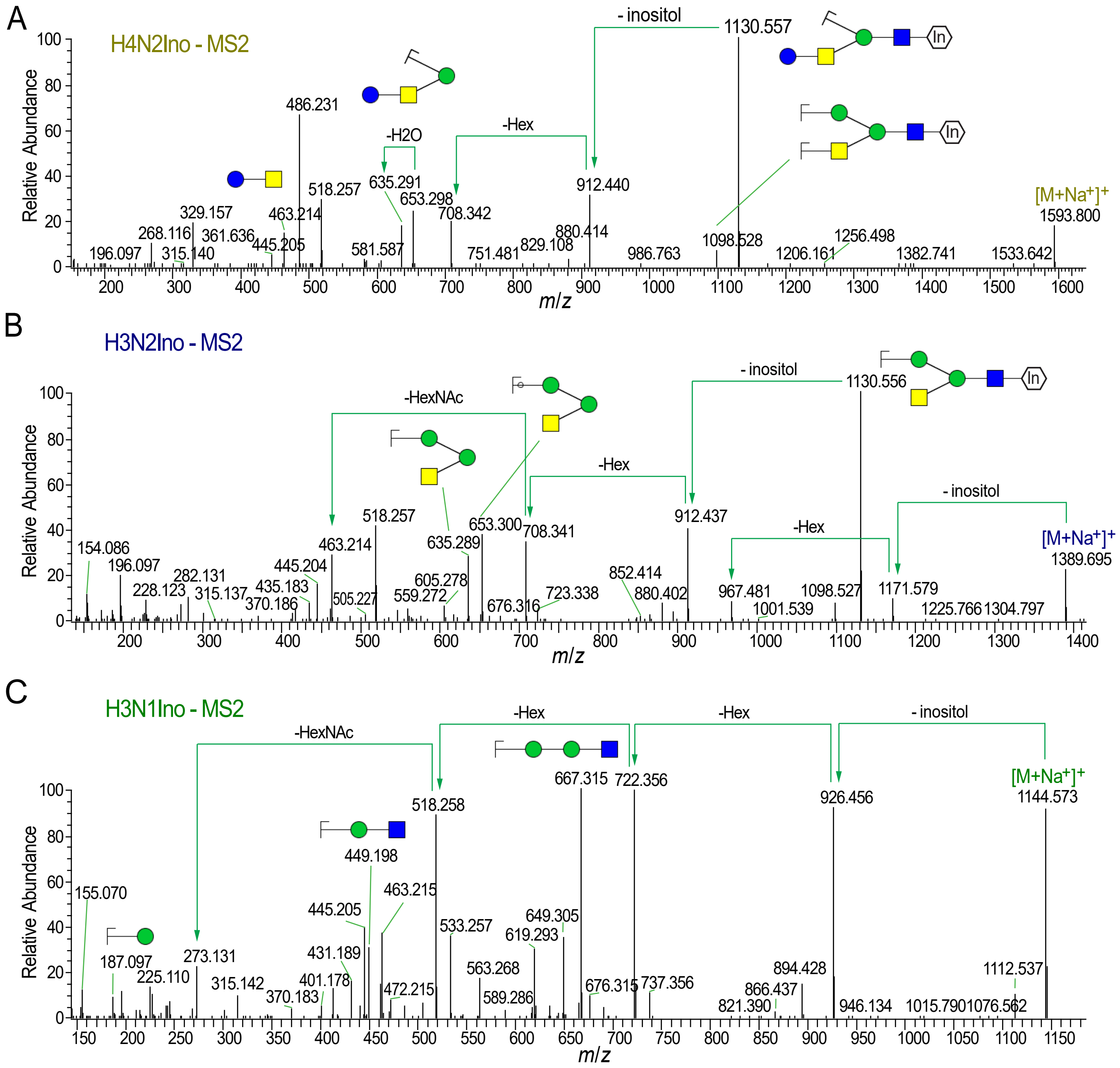

**MALDI-TOF-MS analysis of GPIL preparations**

Fig S6

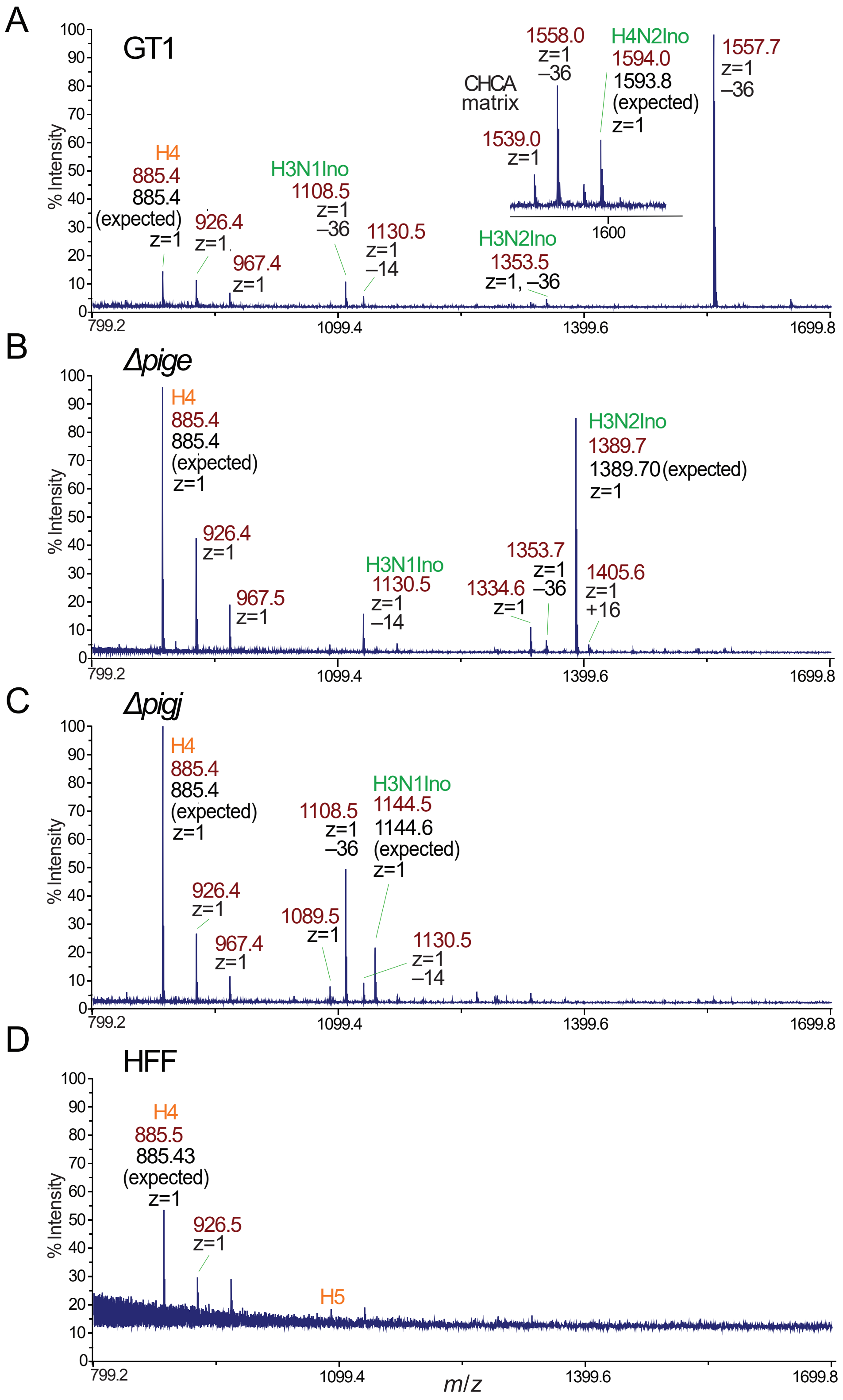

#### nLC-MS analysis of GPIL preparations

Fig S7

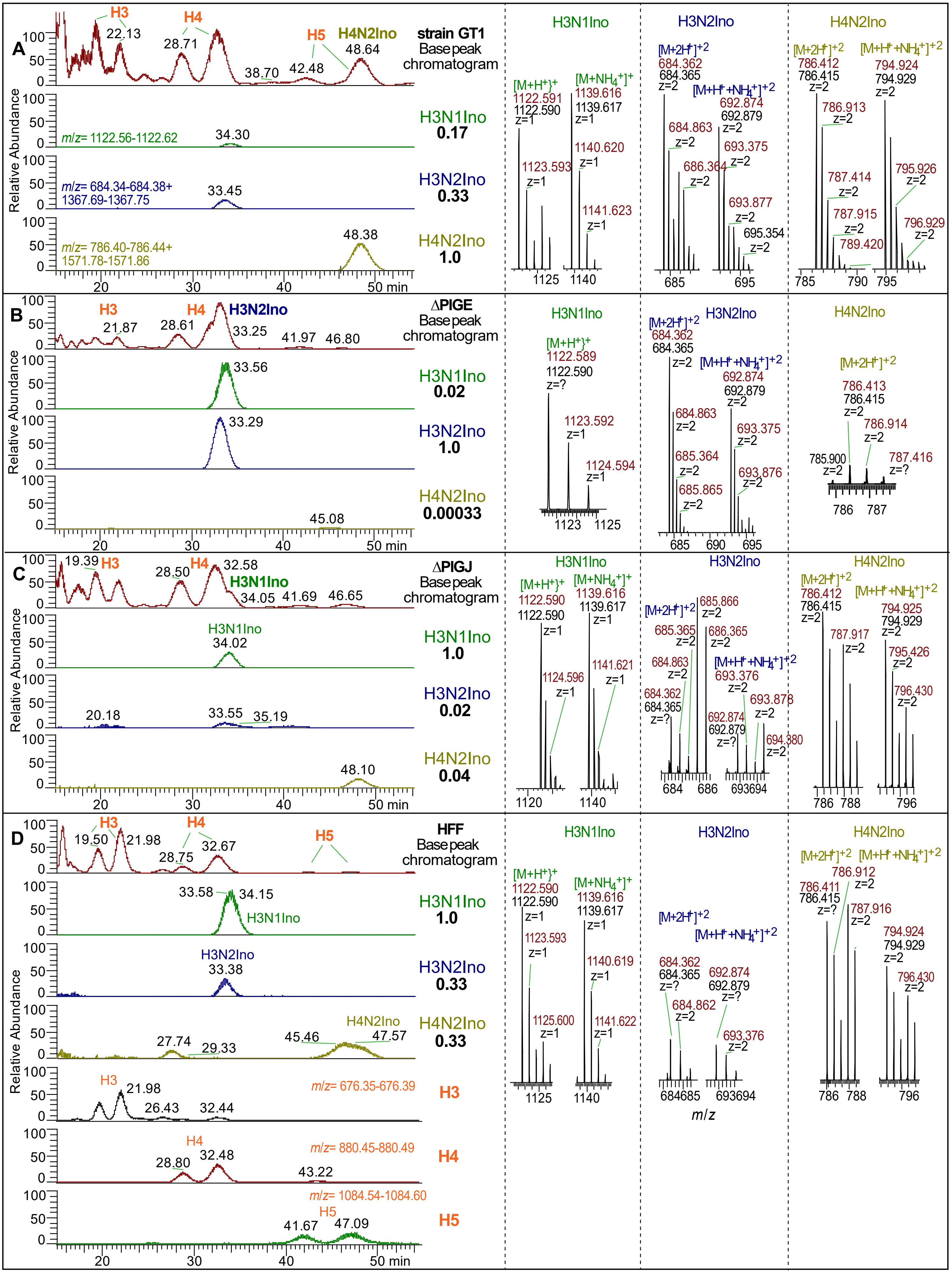

Fig S8

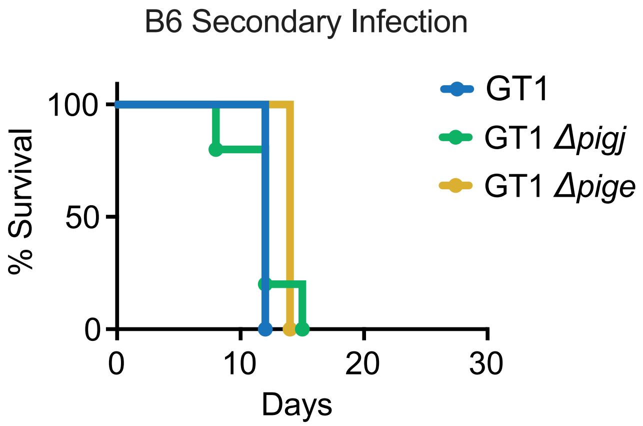
B6 Secondary Infection
% Survival
100
50
0
0
10
20
30
Days
GT1
GT1 Δpigj
GT1 Δpige

Fig S9

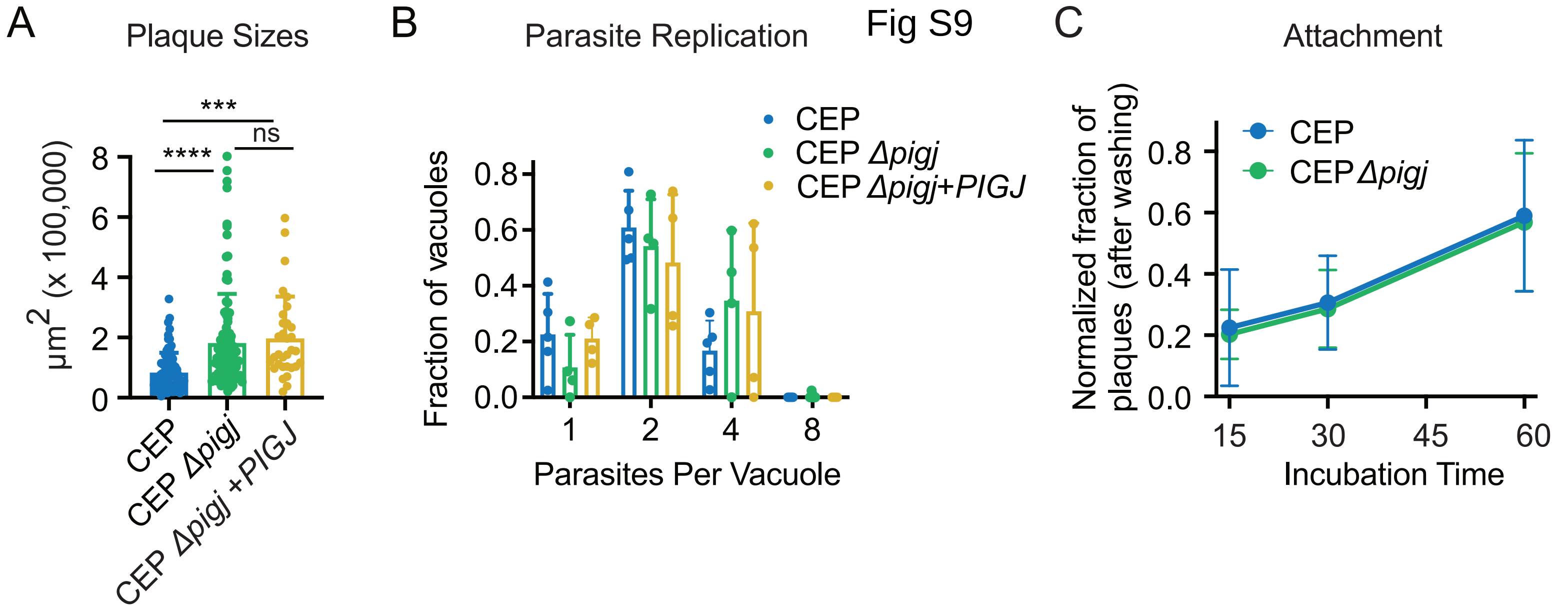

A
Plaque Sizes
μm2 (x 100,000)
8
6
4
2
0
***
****
ns
CEP
CEP Δpigj
CEP Δpigj +PIGJ
B
Parasite Replication
Fraction of vacuoles
0.8
0.6
0.4
0.2
0.0
1
2
4
8
Parasites Per Vacuole
CEP
CEP Δpigj
CEP Δpigj+PIGJ
C
Attachment
Normalized fraction of plaques (after washing)
0.8
0.6
0.4
0.2
0.0
15
30
45
60
Incubation Time
CEP
CEP Δpigj

### Fig S10

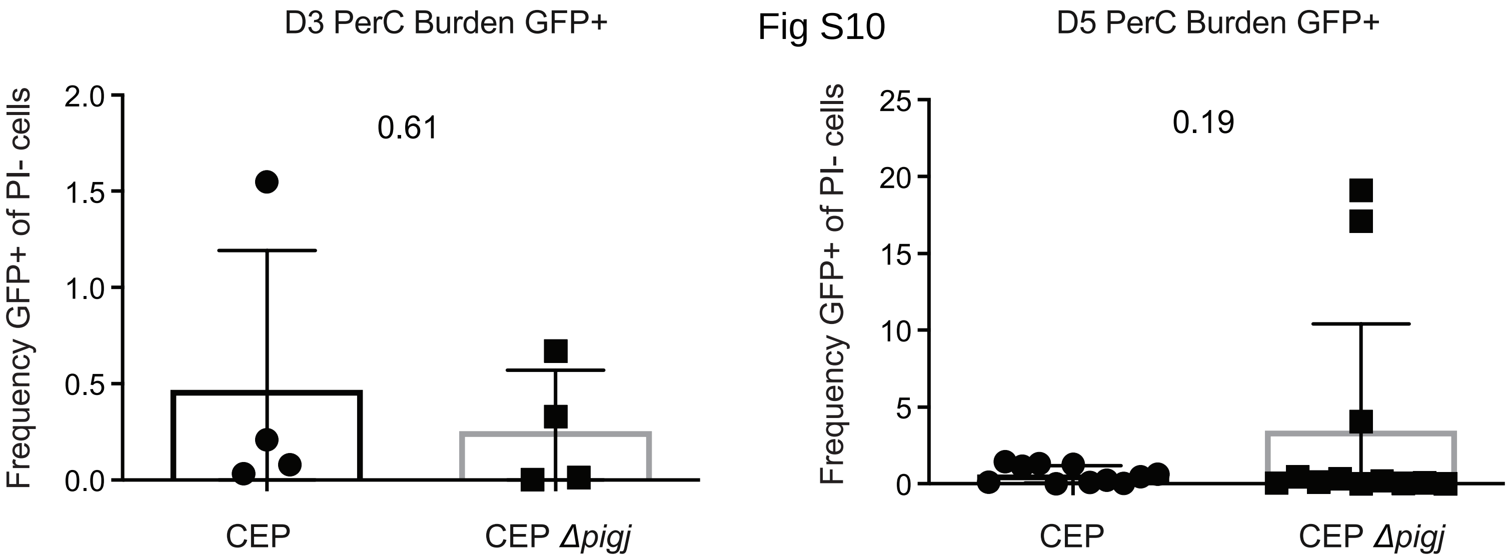
D3 PerC Burden GFP+
Frequency GFP+ of PI- cells
2.0
1.5
1.0
0.5
0.0
0.61
CEP
CEP Δpigj
D5 PerC Burden GFP+
Frequency GFP+ of PI- cells
25
20
15
10
5
0
0.19
CEP
CEP Δpigj

Fig S11

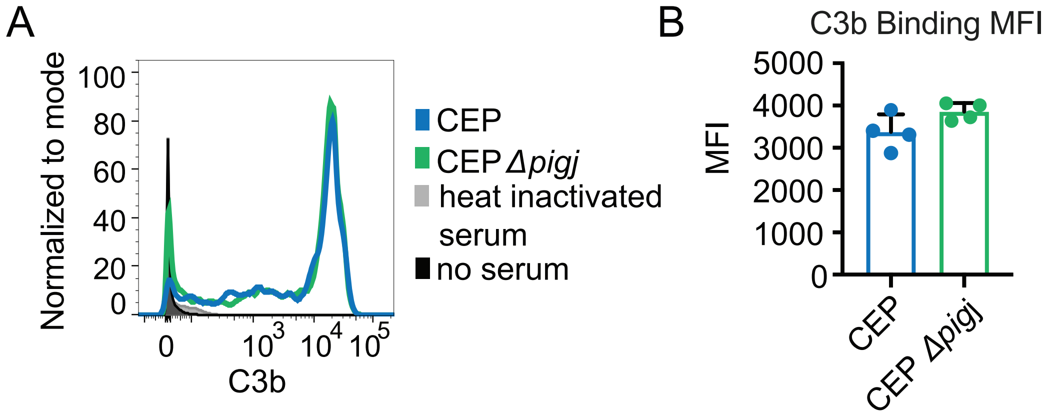
A
Normalized to mode
100
80
60
40
20
0
0
10^3
10^4
10^5
C3b
CEP
CEP Δpigj
heat inactivated serum
no serum
B
C3b Binding MFI
MFI
5000
4000
3000
2000
1000
0
CEP
CEP Δpigj

Fig S12

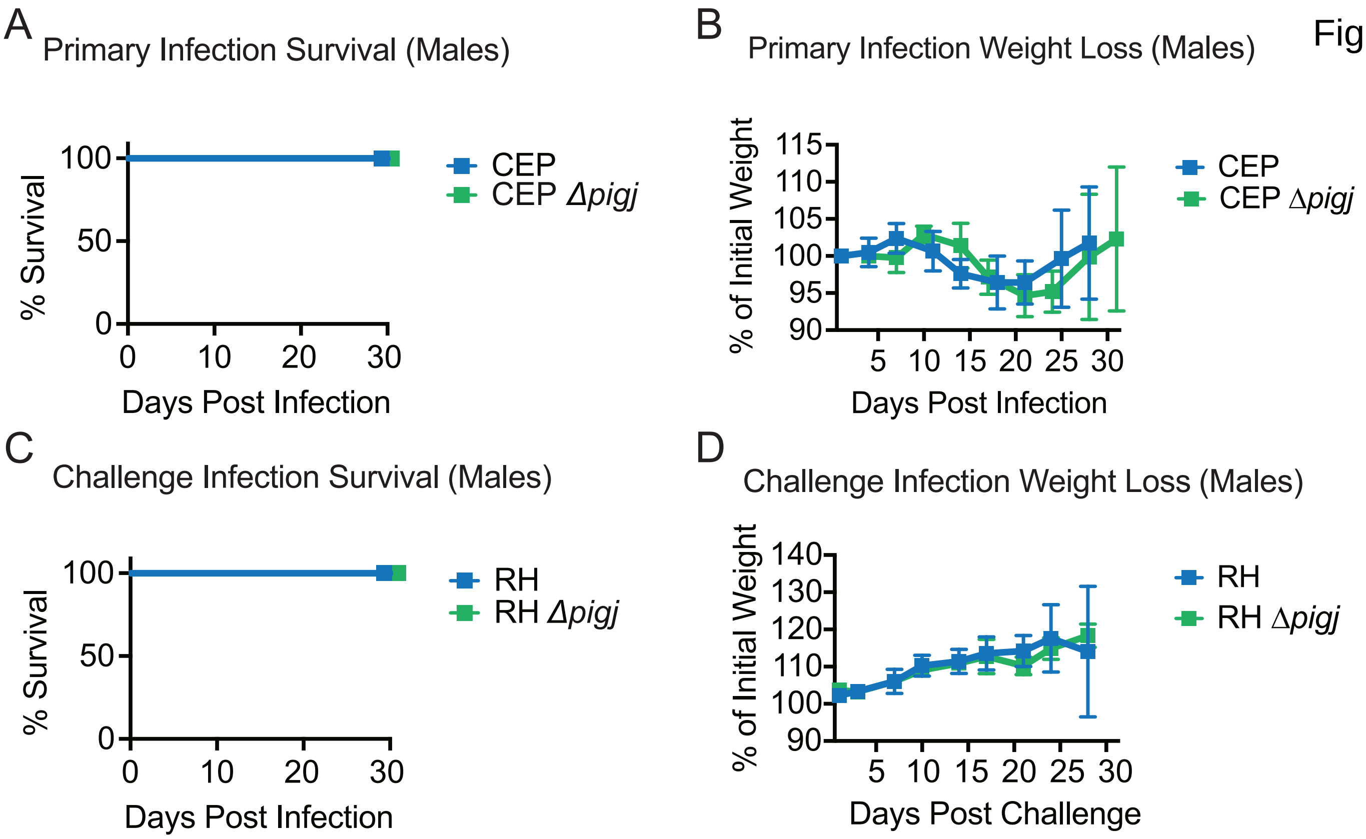
A
Primary Infection Survival (Males)
% Survival
100
50
0
0
10
20
30
Days Post Infection
CEP
CEP Δpigj
B
Primary Infection Weight Loss (Males)
% of Initial Weight
115
110
105
100
95
90
5
10
15
20
25
30
Days Post Infection
CEP
CEP Δpigj
C
Challenge Infection Survival (Males)
% Survival
100
50
0
0
10
20
30
Days Post Infection
RH
RH Δpigj
D
Challenge Infection Weight Loss (Males)
% of Initial Weight
140
130
120
110
100
90
5
10
15
20
25
30
Days Post Challenge
RH
RH Δpigj

### Fig S13

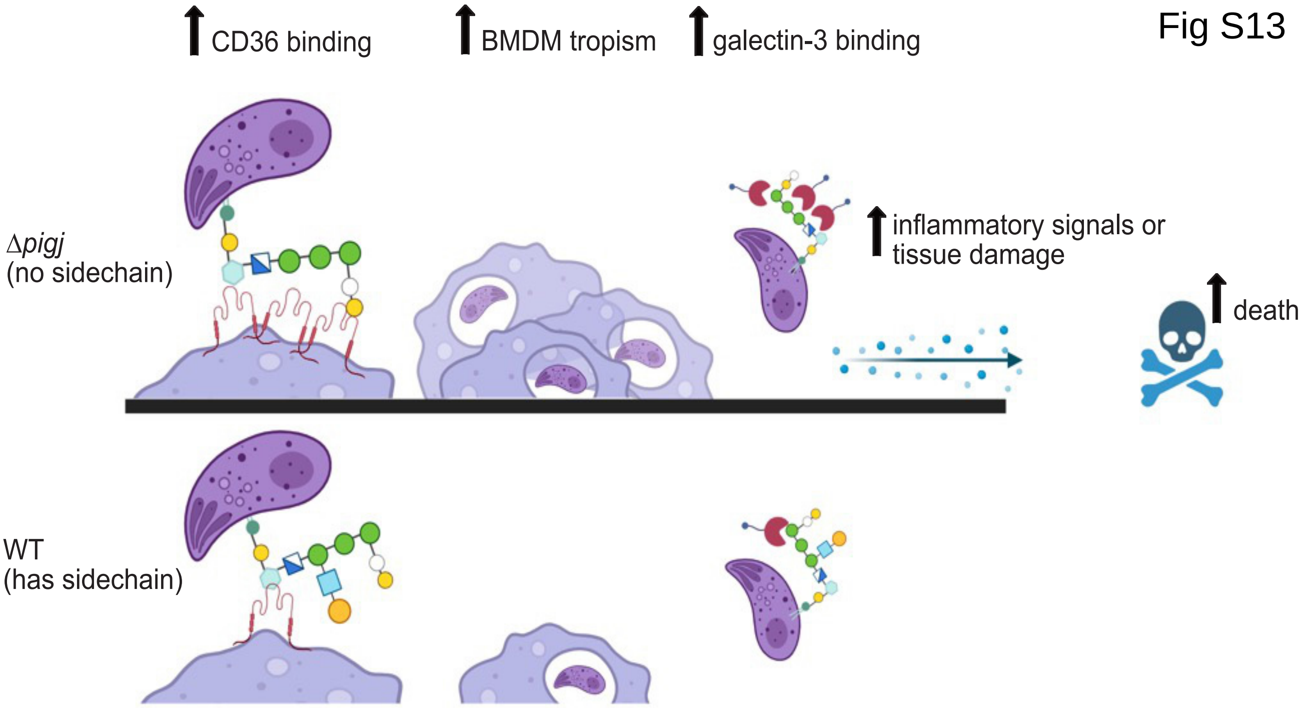
CD36 binding
BMDM tropism
galectin-3 binding
Δ*pigj*
(no sidechain)
inflammatory signals or
tissue damage
death
WT
(has sidechain)
